# Astrocytic cell adhesion genes linked to schizophrenia correlate with synaptic programs in neurons

**DOI:** 10.1101/2021.09.11.459765

**Authors:** Olli Pietiläinen, Aditi Trehan, Daniel Meyer, Jana Mitchell, Matthew Tegtmeyer, Vera Valakh, Hilena Gebre, Theresa Chen, Emilia Vartiainen, Samouil L. Farhi, Kevin Eggan, Steven A. McCarroll, Ralda Nehme

## Abstract

The maturation of neurons and the development of synapses – while emblematic of neurons – also rely on interactions with astrocytes and other glia. To study the role of glia-neuron interactions, we analyzed the transcriptomes of human pluripotent stem cell (hPSC)-derived neurons, from a total of 80 human donors, that were cultured with or without contact with glial cells. We found that the presence of astrocytes enhanced synaptic gene-expression programs in neurons. These changes in neuronal synaptic gene expression correlated with increased expression in the co-cultured glia of genes that encode synaptic cell adhesion molecules, and they were greatly enhanced in the glia in coculture. Both the neuronal and astrocyte gene-expression programs were enriched for genes that are linked to schizophrenia risk. Physical contact between the two cell types was required for the induction of synaptic programs in neurons. Our results suggest that astrocyte-expressed genes with synaptic functions are associated with stronger expression of synaptic genetic programs in neurons and suggest a potential role for astrocyte-neuron interactions in schizophrenia.

## Introduction

Schizophrenia is a severe brain disorder characterized by delusions and hallucinations, impairments in executive function and other cognitive function, and flattened motivation, emotion, and interest^1^. Schizophrenia affects 1% of people globally, but the biological mechanisms underlying the disorder are unknown^2^. Inheritance is a major risk factor for schizophrenia, and recent genetic discoveries have highlighted the quantitative enrichment of genes that are highly expressed by neurons and encode proteins that function at synapses^3–5^. These fundamental aspects of neuronal biology are dependent on interactions with glial cells including astrocytes^6,7^, and raise the question whether cell-nonautonomous effects of glial cells on neurons are relevant to brain disorders such as schizophrenia.

Astrocytes provide neurons with homeostatic support and regulate neuronal development and maturation ^6^. They participate in the formation and shaping of the neuronal network, by regulating synapse generation and elimination, transmission, and plasticity^7–9^. Astrocytes surround neuronal cell bodies and synapses and interact with neurons through a range of contact-dependent and secreted signals that contribute to neuronal maturation^8,10,11^. However, while glial cells are necessary for the functional maturation of neurons^12–15^, many gaps remain in our understanding of the specific cellular and molecular programs that mediate these processes. To investigate the molecular pathways underlying glia-induced neuronal maturation, we performed RNA sequencing of human pluripotent stem cell (hPSC) derived excitatory neurons that were cocultured with mouse glial cells. We reasoned that cellular responses to regulatory interactions between neurons and glia would be in part mediated through mutual changes in cell states that could be detected as correlated gene expression in the two cell types^16,17^. The cross-species cell culture enabled us to separate transcripts by their origin and, therefore, identify molecular dependencies between cocultured neurons and glial cells by studying joint variability in their transcript abundances. Our analysis revealed that astrocyte-expressed synaptic cell adhesion molecules, including *NRXN1*, with established roles in schizophrenia, were associated with induction of synaptic genetic programs in cocultured neurons. We further found that these genes were induced in the glial cells upon coculture with neurons, and that physical contact with neurons was required to induce many of the pro-synaptic effects. These data suggest that the cellular processes in glia that are associated with neuronal maturation involving synaptic programs *in vitro* are relevant to schizophrenia and provide insight into the potential role of astrocytes in psychiatric disorders.

## Results

### Transcriptional profiling of hPSC-derived neurons grown with or without glial cells

To study molecular pathways in glia-neuron interactions, we first assembled a sample set of 32 karyotypically normal hPSC lines (29 iPSC and three ESC lines) derived from neurotypical donors ^15,18^ **Figure 1, Table S1**). These lines were differentiated into excitatory neurons using a well-characterized protocol that combines Ngn2 expression with forebrain patterning factors to robustly generate a homogenous population of excitatory neurons (N= 32 cell lines)^15^. At day four, we plated the neuronal cultures onto a monolayer of mouse glial cells that were prepared as described previously ^19^ from the cortex of neonatal mice (P1-P3). We have previously shown that co-culture with glial cells is necessary for functional neuronal maturation, which we assessed by measuring synaptically driven network activity using Multielectrode arrays (MEA)^15^. To study the effect of glia-coculture on neurons, we first compared the cocultured neurons to a set of five hPSC lines (out of 32) that were differentiated in glia-free condition. In the absence of glia, neuronal cells tended to cluster and clump up together, whereas neurons cultured with glia appeared overall healthier and were more evenly distributed on the glial monolayer (**Figure 2A, Figure S1A**). Consistent with previous work, this showed that the glia-coculture is overall beneficial for the neurons. We then collected RNA sequence data from the 33 cultures at day 28 of the experiment, a point at which the neurons cocultured with glia display robust neuronal morphology and electrophysical activity^15^. To explore differences in developmental trajectories, we also performed RNA sequencing (RNA-seq) at day 4 of the differentiation for the same lines (in 34 independent differentiations) prior to glia coculture, when the differentiating cells resembled neuronal progenitor like cells (NPCs), as described previously^15,20^, totaling to 67 RNA libraries (**Table S1**). All 67 differentiations were carried out in three experimental replicates that were (after confirmation that replicates had similar results) treated as a single, combined sample in the analyses described below.

**Figure 1.**
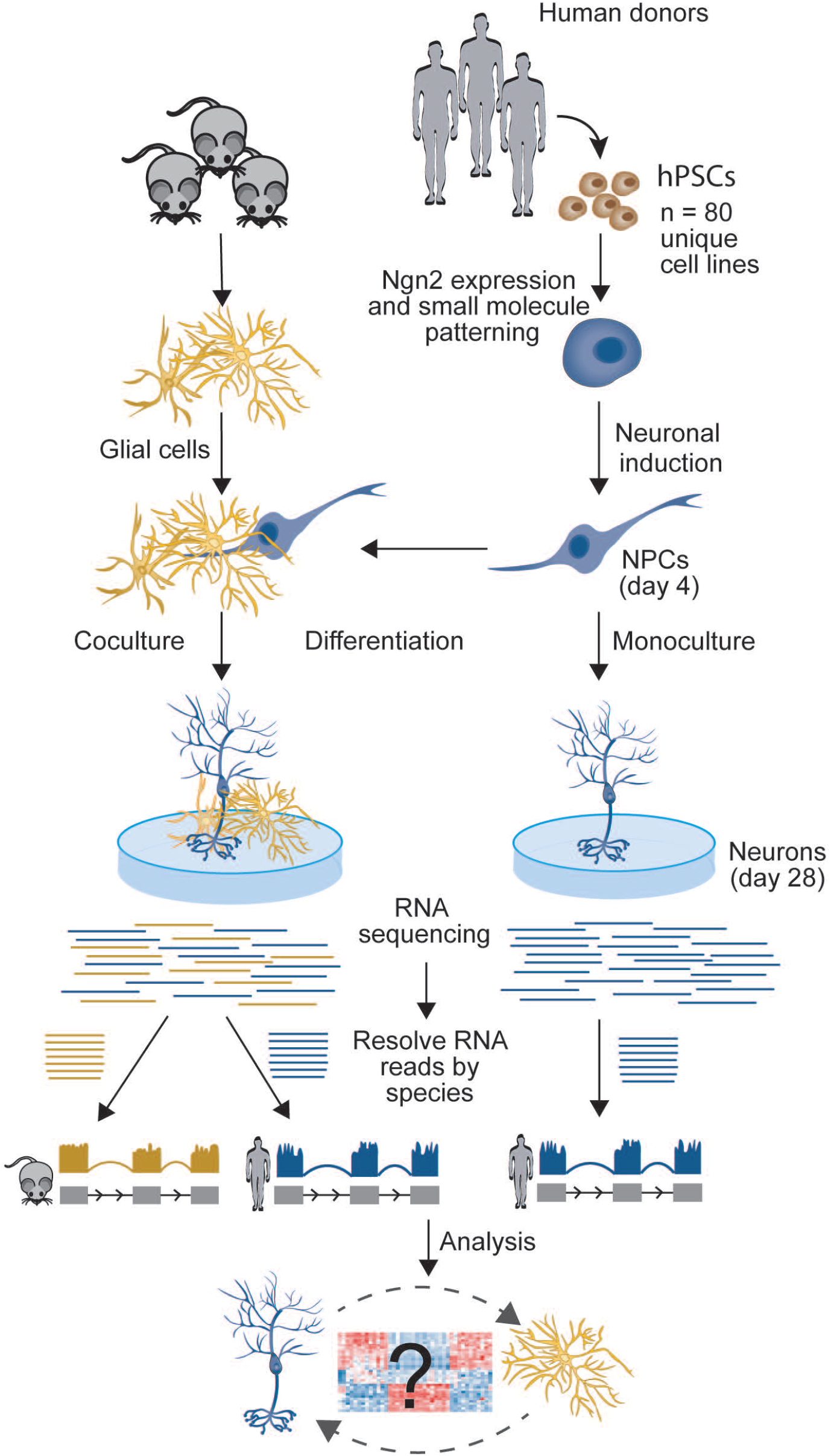
Outline of the experimental design. Neuronal induction with Ngn2 and small molecule patterning was conducted for hPSCs from 80 healthy donors. At day 4 of the differentiation part of the neuronal cultures were plated on mouse glial cells and others were left to differentiate alone. RNAseq was performed at day 28 of the differentiation and the libraries were resolved by species followed by analysis of normalized read counts.

**Figure 2.**
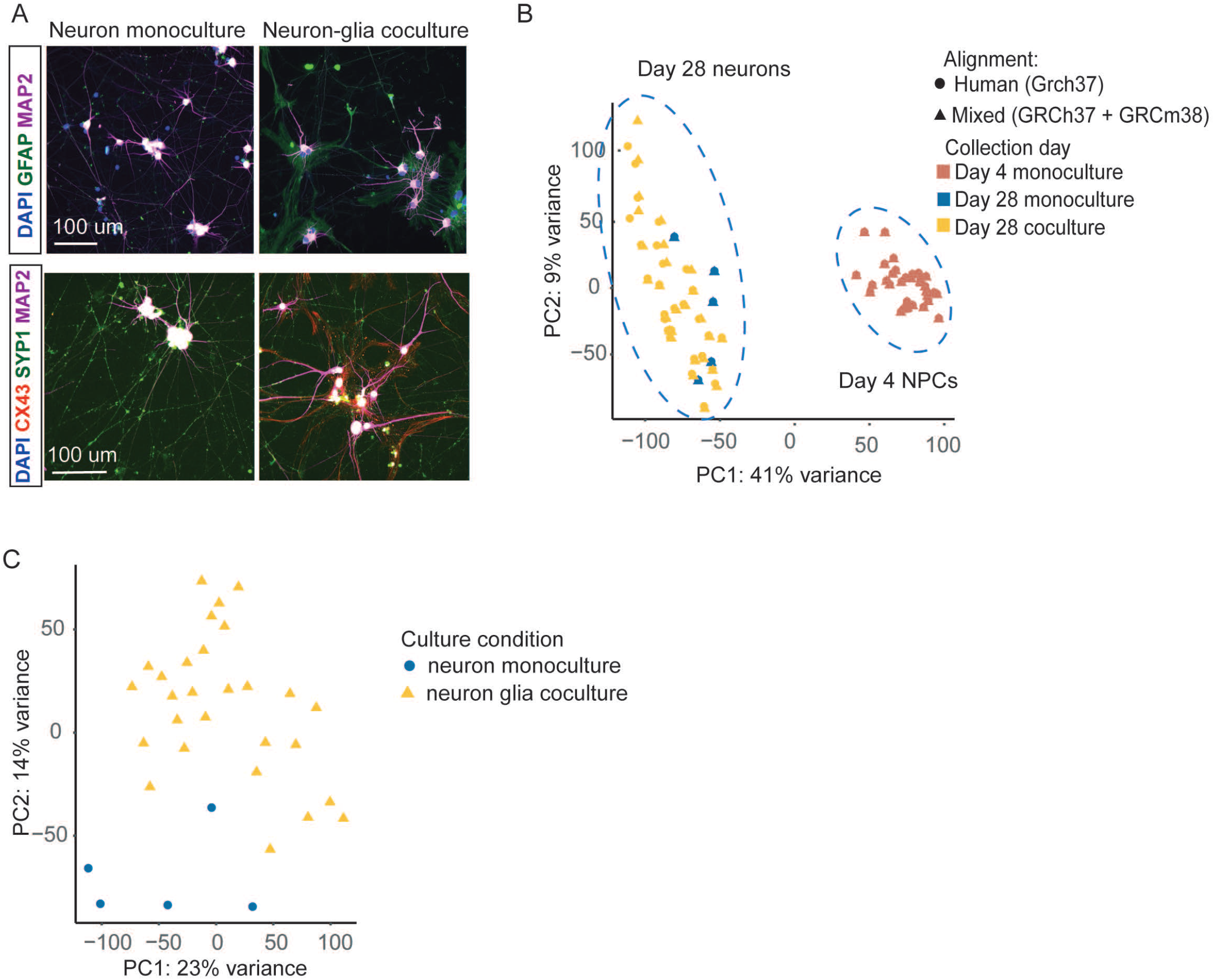
Characterization of of human neurons. A) Representative images of neuron monocultures and neuron-glia co-cultures stained for, top: DAPI (blue) GFAP (green) and MAP2 (magenta), and bottom: DAPI (blue) CX43 (red) Syp1 (green) and MAP2 (magenta), Scale bar= 100um. B) PCA of RNAseq data from day 4 NPCs and day 28 neurons aligned to human and mixed reference genomes. C) PCA of RNAseq data from day 28 neurons aligned to mixed refence genome.

### Leveraging cross-species co-culture to distinguish RNA molecules from different cell types

To characterize cell-type-specific transcriptional effects in the two-species co-culture, we first confirmed that we could reliably distinguish RNA molecules that originated from the human neurons from those that originated from the mouse glial cells^21,22^. To this end, following sequencing, we aligned the reads to a combined Ensembl human-mouse reference genome (GRCh37/hg19 and GRCm38/*mm10*, respectively; GSE63269)^22^ and compared these to read counts obtained after aligning to human genome assembly alone (GRCh37/hg19). We used variance partitioning to estimate the proportion of variability in the gene expression estimates arising from differences between the two alignments. Reassuringly, we found minimal crossspecies mapping (**Figure S1B**). For most genes, the differences in the alignment added little or no variance to the RNA abundance estimates (mean = 0.78 %, median = 0 % of total variance explained). Importantly, only for 329 (of 19,185) genes did the alignment have effects that reached 10% of the inter-sample variance in gene expression (**Figure S1B**). These 329 genes were on average 22.3kb shorter than the rest (95%-CI: 14.1kb-30.6kb p =0.00035, Mann-Whitney) and did not have any specific gene ontology (GO) enrichment (q > 0.05 for all terms). As expected, the alignment to the mixed genome yielded slightly lower (average 3.9%) library size due to reads that mapped ambiguously between the two species’ genomes (average pairwise difference in read counts = 506×10^3^ reads (95%-CI: 765×10^3^ - 247×10^3^), p=0.0002, paired t-test).

A principal component analysis (PCA) of the RNA-expression profiles of the day-28 neurons and day-4 NPCs confirmed that the alignment effect was similar in the coculture and monoculture experiments and did not confound the primary sources of biological variability in the data (**Figure 2B**). Encouragingly, as evident from the superimposed data in PCA, the primary components of gene expression were not affected by the alignment. Instead, the small fraction of genes whose expression estimates were impacted by cross-species mapping were captured in PC7 and PC8 (explaining 5% of the total variance) (**Figure S1C**). Importantly, this was not dependent on the differentiation stage or the presence of glia coculture, suggesting that there was no major bias by experimental condition in the read count estimates. Together, these results confirmed that we could accurately distinguish gene expression in human neurons from that in co-cultured mouse glial cells.

### Neurons grown with mouse glia cells exhibit global changes in gene expression

Having confirmed that we could reliably assign RNA sequence reads to their cell population of origin, we next sought to confirm the neuronal identity of the day 28 neurons (**Figure S1D**). A comparison of the expression of canonical marker genes (**Table S2**) in neurons and day 4 NPCs confirmed the expected temporal reduction of the pluripotency genes *POU5F1*/Oct4 and *MKI67* (Neuron monoculture: β = −1.48, se: 0.24, p = 7.95×10^−9^ and coculture: β = −1.39, se: 0.13, p = 5.4×10^−20^). As expected, neuronal identity genes (*DCX, MAP2, MAPT, NCAM1, RBFOX3, SYN1, TUBB3*) were induced over time in day 28 neurons (both mono and coculture) compared to day 4 NPCs (monoculture: β = 0.36, se: 0.17, p = 0.03; coculture: β = 0.75, se: 0.09, p = 3.1×10^−16^). This was accompanied with modest overall reduction in neuronal progenitor markers genes (*EMX2, HES1, MSI1, NEUROD1, OTX2*; Neuron monoculture: β = −0.46, se: 0.20, p = 0.03; coculture: β = −0.42, se: 0.11, p = 0.0002), suggesting that the neurons might have retained some features of less-mature neurons.

We then focused on characterizing the global transcriptome landscape of the differentiating neurons in both culture conditions. As expected, the largest component of variation in PCA (PC1: 44%) reflected differentiation/maturation time (the difference between the d4 and d28 samples). Interestingly, within the neuron cluster, neurons grown as a monoculture grouped slightly closer to NPCs than neurons cocultured with glia did **(Figure S1E**). This gradual separation became more prominent in PCA after omitting data from the NPCs (**Figure 2C**). Taken together, the data suggested that glia coculture induced global transcriptional changes consistent with greater neuronal maturation. Furthermore, we found that culture-to-culture variability of the cocultured neurons was markedly larger (dispersion: mean = 0.27, median = 0.21) than in neurons in monoculture (dispersions: mean = 0.17, median = 0.11) and in NPCs (dispersions: mean = 0.16, median = 0.12, **Figure S1F**). This suggested that the glial cells introduced an additional source of variance into the neuronal system. We reasoned that this variance could be, at least in part, the result of differences in the glial cell state and composition of the primary mouse cultures and could therefore inform the study of biological drivers underlying glia-neuron interaction.

### Astrocytes are the predominant cell type in the mouse glial cultures

Next, we moved on to characterize the glial cell population in the 28 cocultures by exploring the expression of genes characteristic of specific glia cell types from the mouse (**Table S2**). In line with previous work^19^, we observed predominant expression of genes characteristic of astrocytes (mean: 4.4 log2TPKMs) compared to marker genes for oligodendrocytes (Δ= 2.9 log_2_TPKM, p = 7 ×10^−28^), homeostatic microglia (Δ= 1.0 log_2_TPKM, p =1 ×10^−6^), and microglia (Δ= 3.5 log_2_TPKM, p = 5 ×10^−47^) in the glia coculture (**Figure 3A**).

**Figure 3.**
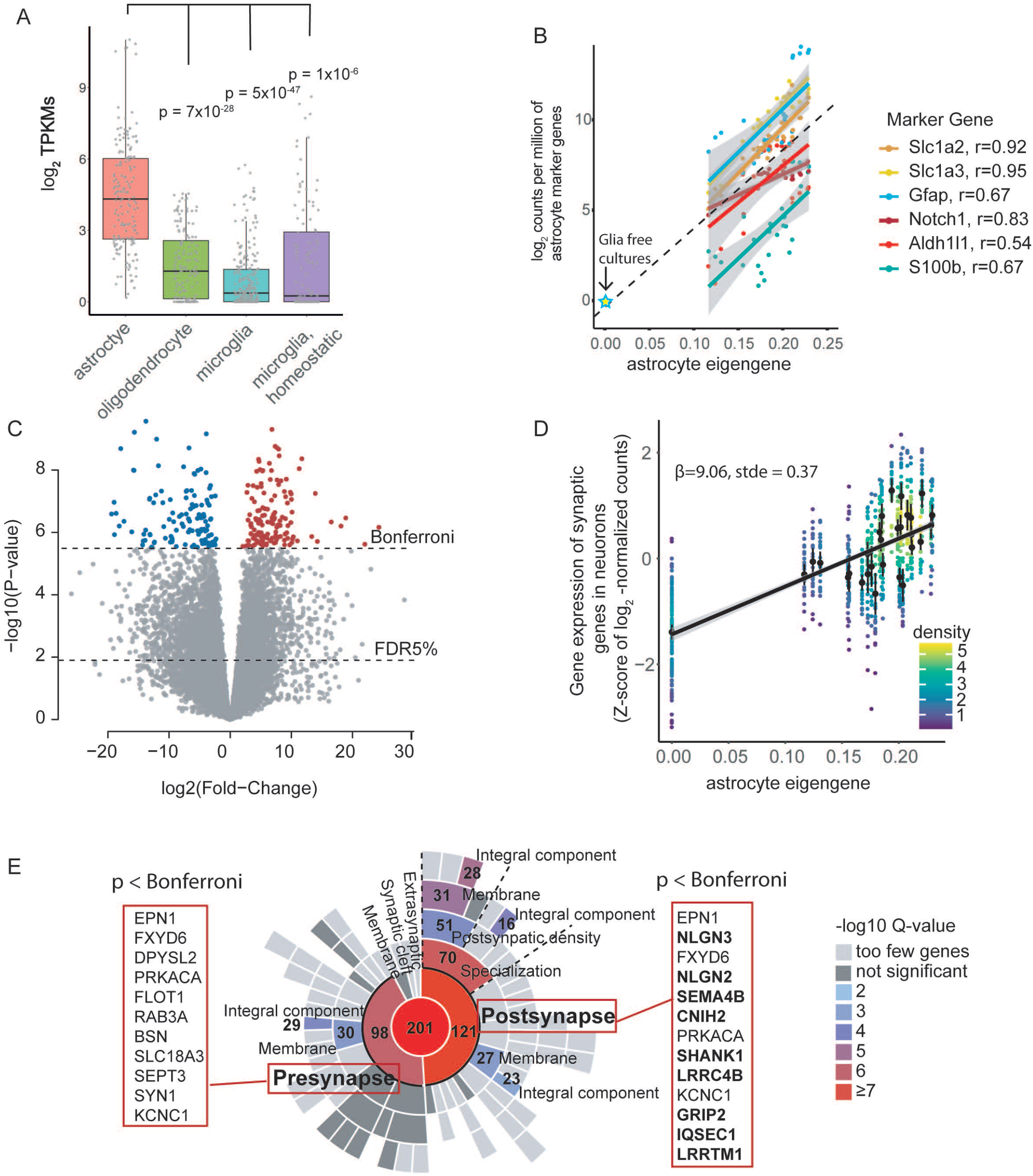
Astrocyte eigengene associates with synaptic gene expression in cocultured neurons. A) Expression of canonical markers for glial cells in the cocultures. B) An astrocyte eigengene was generated by SVD from six astrocyte marker genes. Astrocyte marker genes are highly correlated with the generated eigengene C) Differential expression analysis in neurons of the astrocyte eigengene reveals 4,195 associated genes (1,970 induced, FDR < 5 %), out of which 250 were transcriptome-wide significant (p < 3.0×1ũ-6, Bonferroni, colored genes). D) Synaptic genes were induced in neurons in cocultures with high expression of astrocyte marker genes. E) Astrocyte induced genes in neurons were enriched for synaptic annotations in SynGO.

Guided by our observation of increased variability in cocultured neurons, which we presumed to result from biological variability in glial cell composition and state, we decided to investigate whether the transcriptional profile of the glial cell population in coculture associated with neuronal maturation state. Since astrocytes have been found to regulate neuronal maturation^23^, we specifically sought to determine whether genetic programs related to astrocytes were associated with differences in the maturation. For this purpose, we first estimated the relative abundance of astrocytes in each of the 28 cocultures by combining information from six canonical astrocyte marker genes (*Aldh1l1, Slc1a3, Slc1a2, Gfap, Notch1, S100b*). We used singular value decomposition to generate a single astrocyte eigengene from the marker genes distinctive of astrocytes (**Figure 3B, Table S1**). The marker genes were highly correlated and the eigengene captured 97 % of the variance in the glial cell composition and cell state in the culture. This is consistent with the marker gene transcripts originating from a single shared cell-type^24^ – presumed to be astrocytes – in the glia coculture, and further confirmed the generated eigengene as a relevant proxy for astrocytes in the glial coculture. To measure the full effect of glia-neuron co-culture on gene expression, we included the five glia-free monocultures (out of 32) as the negative control for glia-induced outcomes in neurons. For glia-free cultures the astrocyte eigengene value was set to zero. This was in agreement with an extrapolated value (0.008) from the astrocyte marker gene expression for the glia free cultures (**Figure 3B**). We hypothesized that by leveraging a range of quantitative effects across many cultures we could also better control for the observed qualitative culture differences between the monocultures and cocultures.

### Astrocytes induce transcripts with synaptic functions in neurons

To investigate whether this gene-expression variation in the astrocytes (across the individual cultures) associated with gene-expression variation in the neurons, we used a multifactorial linear model (Limma-voom package^25^) to model each gene’s neuronal expression level, with the astrocyte eigengene and normalized Ngn2 expression as explanatory variables. The analysis revealed transcript abundances of 4,195 human genes (out of 16,694) to be associated (at FDR < 5%) with the astrocyte eigengene value. These included 1,970 positive and 2,225 negative associations to the astrocyte eigengene (**Figure 3C, Table S3**). Overall, the astrocyte eigengene explained 4% — 70% of variance in the 4,195 significantly changed genes (median 21%), while the level of the neuralizing Ngn2 and residual sources of variance had a marked contribution (median: 11% and 67%, respectively) to the gene expression levels as well (**Figure S2A**).

To explore whether the genes whose neuronal expression levels were associated with high astrocyte eigengene values were enriched for specific biological functions, we used gene ontology (GO) enrichment analysis, focusing on 250 genes (135 induced, 115 reduced) that were associated with the coculture astrocyte eigengene at a still-higher-level of significance (p < 3.0×10^−6^, Bonferroni) (**Figure 3C, Table S3**). The analysis of the 135 transcriptome-wide significantly induced neuronal genes revealed most significant enrichments for functions in synapse assembly (GO:0007416, N=9 genes, FC=7.03, q =0.004), axon development (GO:0061564, N=15 genes, FC=3.5, q=0.005), negative regulation of the JAK-STAT cascade (GO:0046426, N=5, FC=13.6, q=0.005), and chemical synaptic signaling (GO:0007268, N=16, FC=3.2, q =0.005) (**Figure S2B, Table S4**). This suggested that astrocytes in the cocultures induced genetic programs related to synapse biology in the neurons. Interestingly, among the most frequently included genes in the enriched categories were genes encoding trans-synaptic cell adhesion proteins including neuroligins (*NGLN2, NGLN3, NECTIN1*), and leucine rich repeat membrane proteins (*LRRC4B, LRRTM1*), with roles in anchoring the synaptic nerve terminals. In comparison, the 115 transcriptome-wide significantly reduced neuronal transcripts in response to astrocytes did not reveal significant enrichment for any specific biological process.

Encouraged by findings from the GO analysis, we next asked whether the larger set of 1,970 induced genes (that passed the FDR < 5 % threshold) were enriched for specific synaptic components in SynGO ^26^. We observed a significant enrichment for genes in the presynaptic component (GO:0045202, N = 98, genes, q = 8.4×10^−7^), and for postsynaptic genes (GO:0098794, N = 121 genes, q=2.9×10^−8^), especially in postsynaptic specialization (N=70 genes, q=3.6×10^−7^), and postsynaptic density (N=51 genes, q=2.1×10^−4^) (**Figure 3D,E, Table S5**). These included 23 transcriptome-wide significantly induced neuronal genes, 13 of which were postsynaptic (**Figure 3D**). Together these results implied that astrocytes in coculture induce or enhance genetic programs in neurons related to pre- and post-synaptic biology, which is consistent with synaptic maturation.

### Replication in isogenic cultures

We next sought to evaluate whether the 1,970 neuron-expressed genes that were correlated with the high astrocyte eigengene values in the accompanying glial cells, were induced in neurons by coculture. We therefore generated pooled cell cultures from an additional 48 donors that were differentiated into neurons either in the presence or absence of glial cells. Neurons from each of the 48 donors were differentiated together in a ‘cell village’ as previously described^27,28^. In brief, neurons from each of the 48 cell lines were induced separately, then mixed in equal numbers 6 days post-induction to form a “village” and differentiated together either with or without glia (**Figure 4A**). At day 28 of the differentiation, we performed droplet-based single cell RNAseq on the neuron villages using the “Dropulation” method, which uses natural donor SNP genotypes as allelic barcodes to assign donor identity for each cell. The pooled experiment enabled the generation of cell cultures from multiple donors that were harmonized for identical culture condition. The cell villages yielded RNAseq data from 76,112 cells with on average 793 cells from each donor and on average 15,586 unique molecular identifiers (UMIs) per cell (**Table S6**). We used t-distributed stochastic neighbor embedding (tSNE) to explore the impact of the glial cell on the state of the cocultured neurons (**Figure 4B,C**). The tSNE demonstrated an even distribution of cells from all donors across the dataset, demonstrating that the cell villages were overall well balanced, and highlighted the reproducibility of the differentiation protocol. Furthermore, we could see that the monocultured neurons clustered separately from the cocultured neurons. This was in line with the PCA from the bulk RNAseq from the discovery cohort. We then constructed meta cells of each donor and compared the 1,970 transcript that were associated with high astrocyte eigengene values between the isogenic neuron villages that were grown with or without the glial cells. Reassuringly, we found that these transcripts had in aggregate higher abundance in the cocultured village than in the glia-free neuron village (difference: 0.21 standard deviations, 95%-CI: 0.22 - 0.19, p = 1.45×10^−43^, t-test, **Figure 4D**). Of the 1,626 genes expressed in both data sets, 65% (1,088) were expressed more highly by neurons in glia-coculture than in neuronal monoculture, with 78% (846) of these differences being nominally significant (FDR < 5%, p=7.0×10^−9^, binomial test) (**Figure 4E, Figure S3A, Table S7**). In summary, these results confirmed that many of the transcripts that were found to associate with high astrocyte eigengene values in the initial experiments were also induced in neurons in glia-cocultures in independent experiments. This is consistent with the idea that astrocyte biology instructs, rather than merely responding to, the neuronal changes.

**Figure 4.**
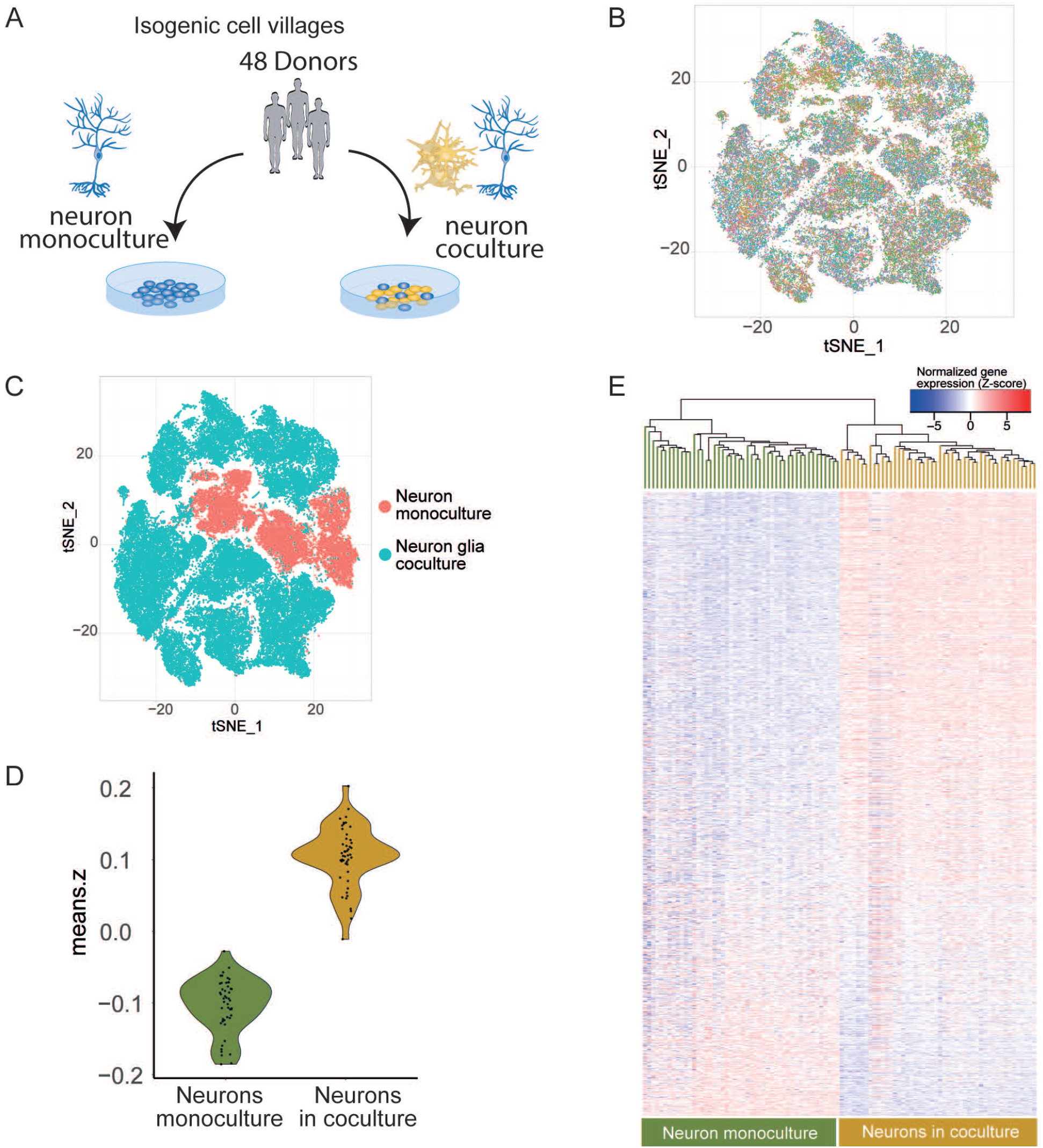
Cell villages of isogenic neurons grown in monoculture or in coculture. A) Study design for single cell RNAseq of pooled neuron villages from 48 donors. B) t-SNE projection of scRNAseq data color-coded by donor. C) tSNE projection of scRNAseq data color-coded of by culture condition (neuron monoculture or coculture with glia). D and E) Neurons in coculture have higher expression of genes associated with the astrocyte eigengene.

### Neurons induce the expression of genes for cholesterol synthesis and synaptic functions in astrocytes

Having established that glia cocultures that have high expression of astrocyte marker genes associate with induction of synaptic gene programs in accompanying neurons, we wondered what biological processes in astrocytes underlie these effects. To address this, we first generated RNAseq data from glial cells in coculture with neurons at day 28 of neuron differentiation (N = 5, 4 replicates each) and from glial cell monocultures of the same batch cultured at the same time (N = 4, **Figure 5A**). The presence of neurons resulted in substantial changes in astrocyte gene expression, with 1,426 (out of 11,205 evaluated) genes differing in expression between the coculture and monoculture conditions (p < 4.5×10^−6^, Bonferroni; 668 induced in coculture with neurons). To investigate which biological processes were affected in the glial cells, we performed a GO enrichment analysis. The enhanced transcripts were particularly enriched for genes with functions in the cholesterol biosynthetic process (GO:0006695, N=15 members, 7.8-times more genes relative to background, q= 6.3×10^−8^, **Figure S4, Table S8**). The induction of the cholesterol synthesis is in line with previous reports suggesting that astrocytes are a major provider of neuronal cholesterol. The most significantly induced genes included *Apoe*, which encodes a lipoprotein that shuttles cholesterol from astrocytes to neurons (log_2_Fold-change=3.4, p=1.8×10^−14^, **Figure 5B**) and Clu (ApoJ) which is also involved in cholesterol shuttling (log_2_Fold-change=1.4, p=7.0×10^−11^). Along with the cholesterol synthesis genes, coculture with neurons enhanced glial expression of many genes with annotated roles in synaptic physiology (GO:0099536, N = 48, 2.4-fold enrichment, q=2.0×10^−6^). These included genes that encode the astrocyte-specific glutamate transporter (Slc1a3/Glast) and the glutamate-ammonia ligase (Glul) that catalyzes the conversion of glutamate, which is taken up from the synapse by astrocytes, to glutamine for recycling back to neurons (**Figure 5B**). These gene-expression changes align with the increase in glutamate-driven activity of the neurons that is observed in glia-coculture and suggest that the glial cells actively participate in the glutamate cycle *in vitro^15^*. Genes encoding synaptic cell adhesion molecules *Nrxn1, Lrrc4*, and *Nlgn3* were also strongly upregulated in glial cells in the presence of neurons (*Nrxn1*: log_2_Fold-change=1.9, p=4.5×10^−12^, *Lrrc4:* log_2_Fold-change=3.2, p=2.3×10^−6^, *Nlgn3:* log_2_Fold-change=1.9, p=8.4×10^−7^, **Figure 5B**). Intriguingly, Nrxn1, Nlgn1 and Nlgn3 have recently been demonstrated to be abundant in astrocytes and have likely roles in the astrocyte processes that surround the synaptic contacts at the tripartite synapse^29,30^. Nlgn1 was also moderately upregulated in the cocultured glial cells (log_2_Fold-change=0.9, p= 2.1×10^−5^). Additionally, we found that Lrrrc4 is also expressed in astrocytes, and induced by neurons. In comparison, the genes whose expression was reduced by neuronal coculture were enriched for roles in developmental processes, cell structure and motility (**Table S9**). Together these results demonstrated that glial cells undergo extensive changes in the cell state in response to neurons and suggested that astrocytes actively participate in neuronal biological processes *in vitro*.

**Figure 5.**
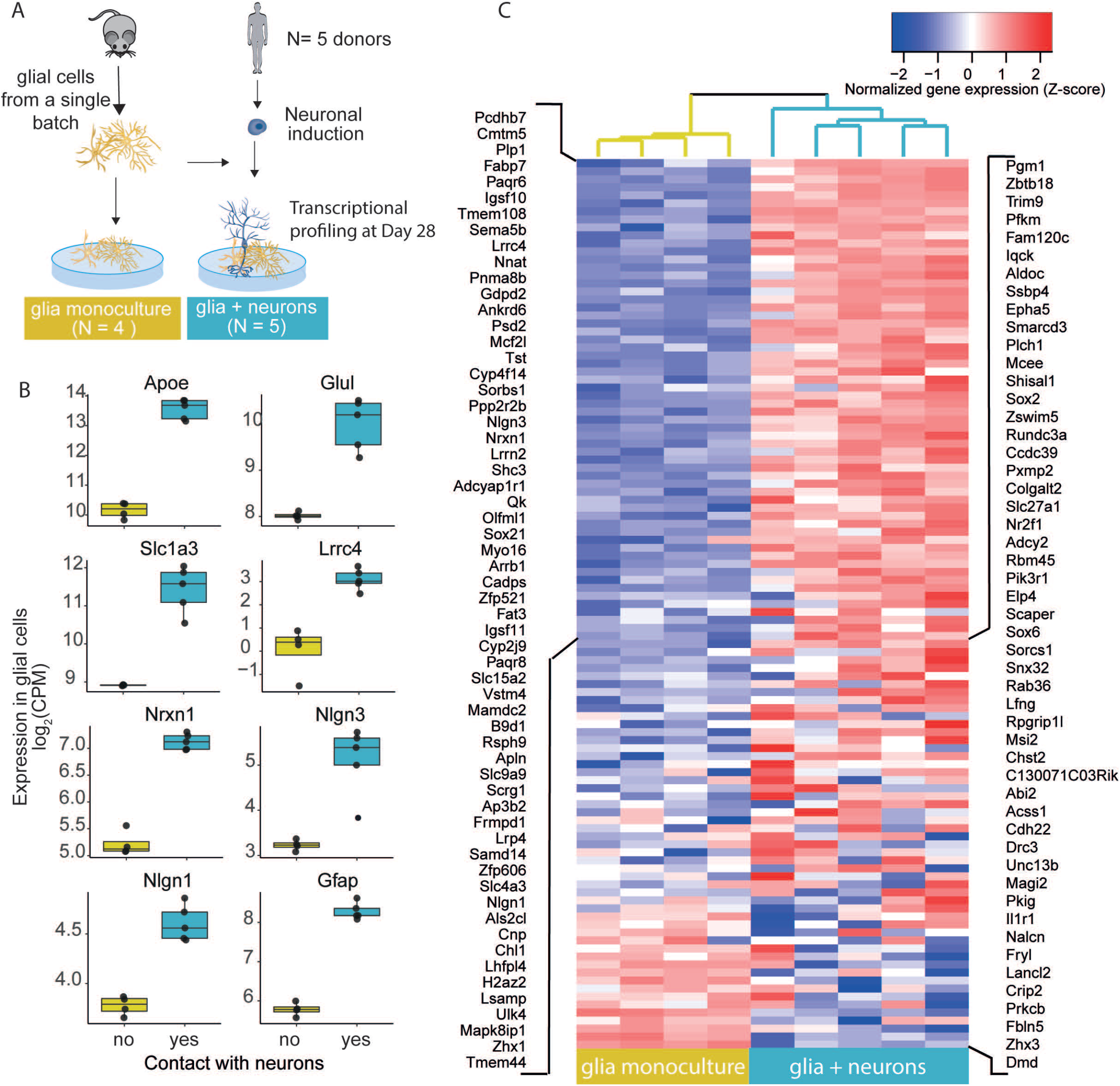
Neuronal impact on glial gene programs. A) Schematic of the experimental design. Gene expression of glial cells was compared in glia monocultures and in coculture with day 28 neurons. B) Neuronal co-culture induced expression of genes in glial cells that were associated with neuronal synaptic maturation. C) Neuron presence induces expression of transcripts of synaptic members including synaptic cell adhesion molecules.

### Astrocyte expressed genes induce synaptic programs in neurons

Encouraged by the observed neuron-induced changes in the glial cells, we asked whether these transcriptional changes in the glial cells were also associated with the synaptic maturation of neurons in coculture. For this purpose, we first condensed the neuronal expression profile of the 1,970 astrocyte-induced genes by singular value decomposition to a single eigengene in the discovery data set (**Table S1**). The generated neuron eigengene values were similar to those of the 135 transcriptome-wide significantly astrocyte-induced genes in neurons (r_median_ = 0.65) and were highly correlated (r=0.68) with the astrocyte eigengene, confirming that the neuron eigengene captured relevant variation in gene expression in the neurons (**Figure 6A**).

**Figure 6.**
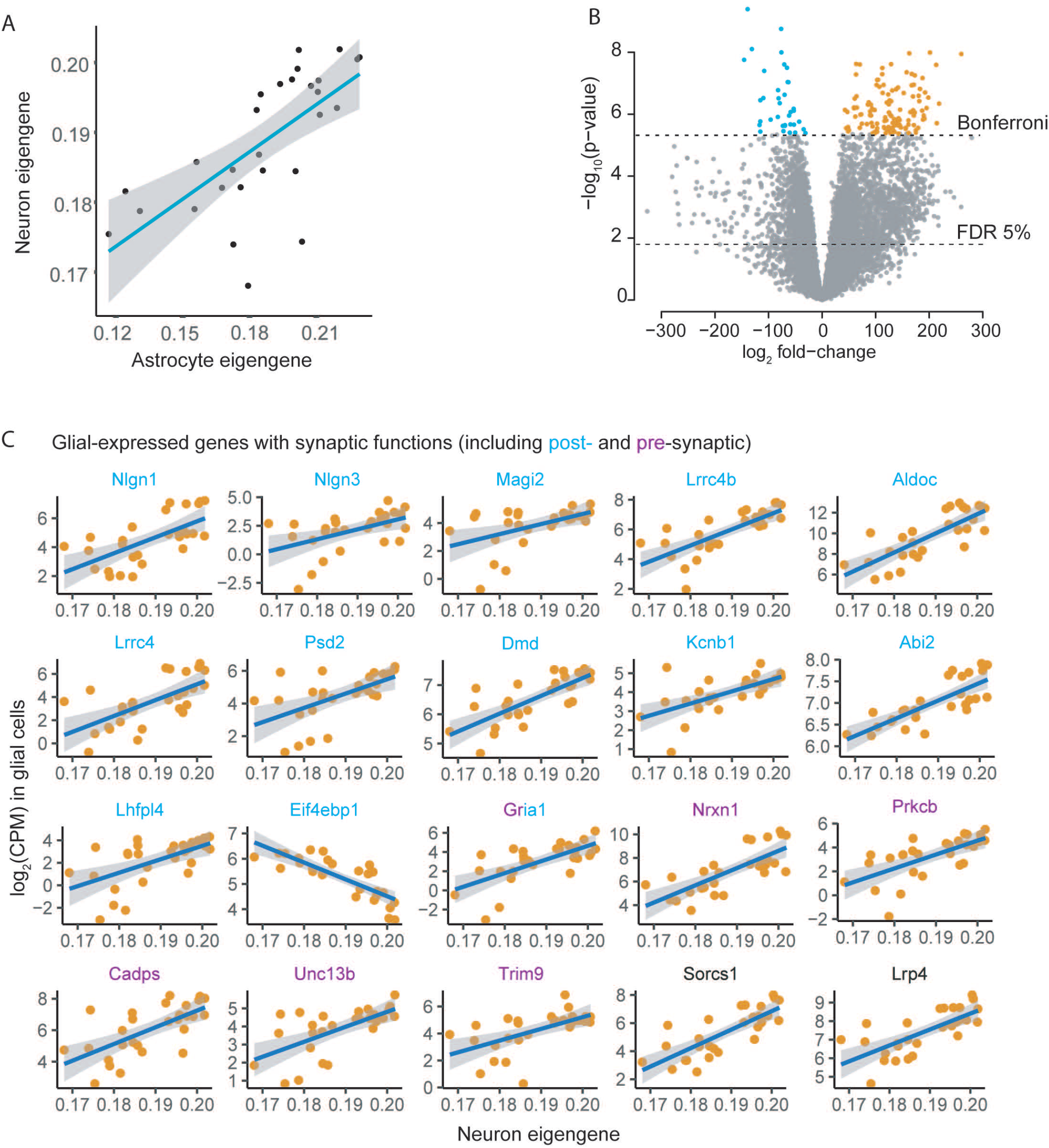
Differential expression of the neuron eigengene in glial cells. A) The neuron eigengene is highly correlated (r=0.66) with expression of astrocyte marker genes (astrocyte eigengene). B) Expression of 159 glial-genes is associated with the neuronal eigengene (4.3×10^−6^, Bonferroni). Differential expression analysis revealed 123 glial genes whose high expression is associated with neuronal maturation in coculture (orange) and 36 genes with negative association to the neuronal maturation state (blue). B) Association of expression of glial genes with neuron eigengene. Postsynaptic genes are labeled in blue, presynaptic genes in purple and other genes are in black.

We then studied the association between the expression levels of individual glial genes (total 11,627 mapped mouse genes) and the maturation state of the cocultured neurons defined by the neuron eigengene value. The analysis of the bulk RNAseq in the discovery sample set revealed transcriptome-wide significant association of transcript levels of 159 glia genes with the neuron eigengene (p < 4.3×10^−4^, Bonferroni). Out of these, 123 genes were positively correlated with the neuron eigengene of the more mature neuronal state and 36 were negatively associated with the neuron eigengene values (**Figure 6B, Table S10**).

To gain insight into the likely source cell type of the 159 glia gene transcripts, we explored their expression in a single cell RNAseq atlas of adult mouse brain^31^. Comparison of the expression patterns of the meta cells from the brain atlas revealed that the 123 genes, whose high transcript levels in glia was associated with higher expression of neuron eigengene, were predominantly expressed by astrocytes compared to other glia cell types (p = 6.9 ×10^−26^, t-test) (**Figure S3B**). In contrast, the 36 negatively associated glia transcripts showed no evidence of preferential expression between the glial cell types in the mouse brain (p=0.81). Moreover, the 36 transcripts had significantly reduced expression in mouse brain astrocytes compared to the 123 genes that were associated to induced neuron eigengene values (p=0.0035, t-test, **Figure S3B**). These results demonstrated that the individual genes associated with synaptic programs in neurons were abundantly expressed by astrocytes in the mouse brain and suggested a role in pathways related to astrocyte-neuron interaction.

### Expression of synaptic cell adhesion molecules by glial cells associate with synaptic programs in neurons

We next asked whether these 159 glia genes reflected specific neurobiological processes that could underlie the interactions with neurons. GO term analysis of the 159 glia transcripts revealed an enrichment of genes with synaptic membrane functions (GO:0097060, N = 13 genes, FC = 4.4, q = 0.002) (**Table S11**). Importantly, this enrichment was driven by genes whose high expression in glial cells induced high expression of synaptic genes in neurons. Reduced expression of only one pro-synaptic gene in glial cells, *Eif4ebp1*, encoding for Eukaryotic translation initiation factor 4e was associated with high neuron eigengene values. Encouraged by these results, we explored curated synaptic annotations from SynGO to confirm 19 glia-expressed genes with annotation to synaptic component, out of which 13 were postsynaptic (q = 0.007) and 6 presynaptic members (q = 0.25) (**Figure 6C, Figure S3C, Table S12**). Remarkably, we noticed that many of these genes were synaptic cell adhesion molecules including *Nrxn1, Nlgn1, Nlgn3, Lrrc4, and Lrrc4b*, similarly to what we found was induced in the glial cells after contact with neurons. Further evidence for synaptic cell adhesion molecules came from the synaptic scaffold proteins Magi2 and Sorcs1 which associate with neuroligins and Nrxn1, respectively, in the synapse^32,33^.

Astrocytes secrete many molecules, including neurotransmitters, modulators, and trophic factors, via regulated exocytosis of synaptic-like vesicles^34^. Interestingly, several of the identified presynaptic genes in the glia have annotated roles in regulating the synaptic vesicle cycle in SynGO (Cadps, Nrxn1, Unc13B, and Prkcb), suggesting a potential additional role for these genes in regulating the secretory vesicle cycle in glial cells. In addition, among the most significantly associated genes was the gene encoding the astrocytic low density lipoprotein receptor related protein Lrp4 (**Figure 6C**), which has roles in regulating glutamate transmission^35^. Together these results suggested that genes encoding proteins with both pre- and post-synaptic functions and members of the tripartite synapse are expressed in glia cells and associated with induced expression of synaptic genetic programs in cocultured neurons.

In comparison, the astrocyte genes negatively associated with the neuronal eigengene did not reveal significant enrichment for specific biological processes. Interestingly, however, the most significantly reduced gene encoded for interleukin 6 family neuropoietic cytokine Lif (leukemia inhibitory factor). Among its many roles, Lif promotes self-renewal and neurogenesis in neuronal stem cells, as well progression of astrocyte precursor cells to mature GFAP^+^ astrocytes ^36–38^. Therefore, the reduced Lif expression could indicate progression from neuro-/glio-genesis to more-mature cell states. In line with this, *Gfap* was found significantly induced in the glial cells by coculture with neurons (log_2_ fold-change =2.5, p=7.9×10^−14^, **Figure 5B**). Furthermore, high expression of *Cmmt5*, a marker of late astrocytes^39^, was the gene most significantly positively associated to the neuronal eigengene. These results suggest that the maturing hPSC-derived neurons enhance the maturation of the accompanying glial cells.

### Neurons stimulate pro-synaptic gene expression programs in glial cells

After identifying the set of 123 astrocyte-expressed genes whose high expression correlated with the maturation of the accompanying neurons, we examined whether these genes overlapped with those that were induced in the glial cells after contact with neurons. Out of the 123 genes, 111 were reliably detected in the glial cell monocultures, and 68 had significantly higher expression (adjusted p-value < 0.05) in glial cells in the presence of neurons (**Figure 5C**, **Table S13, Figure S5C**). This overlap was highly unlikely to have occurred by chance (p = 5.5×10^−19^, binomial test). Of the 18 identified synaptic genes (**Figure 6C**), eight were induced in astrocytes in the presence of neurons (**Figure 5C**). These included all except for one of the synaptic cell adhesion molecules (*Nrxn1, Nlgn1, Nlgn2, Lrrc4*). Similarly, we found that out of the 36 transcripts whose low expression in the glia-coculture was associated with neuronal maturation, 16 were reduced in expression in the glial cells in the presence of neurons (**Figure S3D**). Taken together, our data suggested that glial cells undergo adaptive changes in response to differentiating neurons. These include induced levels of transcripts of synaptic cell adhesion molecules, whose expression in glial cells is in turn associated with advanced maturation of accompanying neurons in coculture.

### Physical contact with astrocytes promotes the induction of synaptic gene-programs in neurons

The association of neuronal-synaptic gene expression with astrocyte expression of synaptic cell adhesion molecules raised the question of whether physical contact between neurons and glial cells was required for the induction of the synaptic gene expression programs in the neurons. To evaluate this, we carried out experiments (with five neuronal lines) in which neurons were differentiated either in a glia-coculture or in a glia-sandwich culture (**Figure 7A**). The sandwich culture enables the exchange of soluble factors in the culture media between the two cell types but prevents any physical contact. On the other hand, in coculture the cells are free to interact by both physical contact and by exchanging soluble factors. We reasoned that differences in the transcriptome between neurons in the two culture conditions would indicate effects brought about by physical contact between glial cells and neurons. We generated bulk RNAseq data at day 28 of the differentiation from the five lines in the two conditions (4-3 replicates each, N = 35 samples). A PCA and hierarchical clustering of the transcriptomic data clustered the isogenic neuron lines by culture condition into those that were differentiated in coculture and those in sandwich culture (**Figure 7B**, **Figure S5A**). This suggested that physical contact between glial cells and neurons was able to induce reproducible changes in neuron state across multiple isogenic experiments.

**Figure 7.**
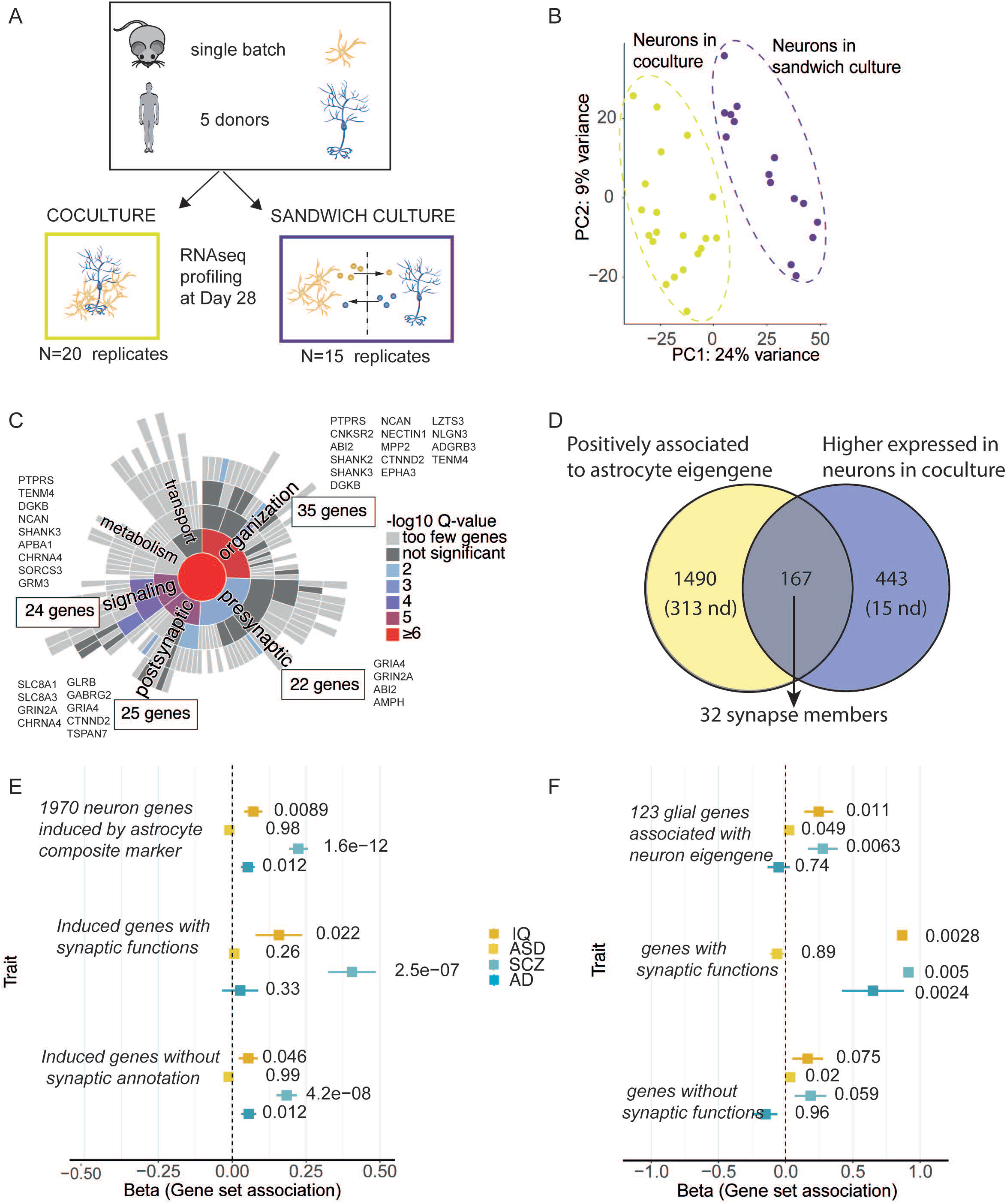
Synaptic programs in neurons are enhanced by physical contact with glial cells and involve genes that associate to schizophrenia. A) Experimental design for isogenic sandwich and coculture experiments. B) PCA of transcriptome of neurons in coculture and sandwich culture. C) Physical contact with glial cells enhances exoression of synaptic genes in neurons. Genes associated with the astrocyte eigengene are highlighted (N=83). D) Overlap of induced genes in coculture and genes with positive association to the astrocyte eigengene. 10% of genes induced in coculutre are associated with high astrocyte eigengene values, nd= not detected. E and F) Gene-set association analysis for four central nervous system traits in neurons (E) and glial cells (F). P-values for gene-set associations for each trait and gene set are indicated in the plot. Astrocytes induce transcripts encoded by genes associated with schizophrenia in neurons (E), and glia-expressed genes that promote neuronal maturation associate with schizophrenia (F).

We next performed a differential gene expression analysis to identify genes whose expression levels were changed in neurons by physical contact with glial cells. The analysis revealed 1,691 genes which significantly (adjusted p-value < 0.05) differed in expression between neurons in coculture and in sandwich culture (**Figure S5B**, **Table S14**). Out of these, 625 were more highly expressed in the cocultured neurons (adjusted p-value < 0.05). We then used GO-term enrichment analysis to investigate whether specific biological processes would be differentially affected in the neurons in the two conditions. The 625 genes with higher expression in glia-cocultured neurons (with glial contact) were enriched for components of nervous system processes and functions in the synapse, with the strongest individual enrichment for chemical synaptic transmission (GO:0007268, N = 61 genes, FC = 2.7, q = 1.6×10^−9^, **Table S15**). A systematic analysis for curated synaptic components in the SynGO database confirmed 104 synaptic genes (q = 4.2×10^−11^), 83 with known roles in synapse processes (q = 1.7×10^−6^), in synaptic organization (N = 35, q= 1.9×10^−5^) and signaling (N=24, q =1.9×10^−5^) (**Figure 7C, table S16**). This suggested that the physical contact between the two cell types upon coculture enhanced synaptic gene programs in neurons beyond effects resulting from soluble factors.

We next asked whether the neuronal genetic programs that were associated with high astrocyte eigengene values could be in part driven by contact-dependent mechanisms with glial cells. A comparison of expression between isogenic neurons in sandwich culture and coculture revealed on average higher expression of the 1,970 genes (1,657 detected) that were associated with the astrocyte eigengene (p =1.9×10^−28^, paired t-test; **Figure S5C,D**). We next identified the overlapping set of genes between the 625 genes with higher expression in coculture than in sandwich culture and the 1,970 genes that were associated with astrocyte eigengene values. Intriguingly, we found 167 genes that had both high expression in cocultured neurons and were induced by high astrocyte eigengene values. This overlap was significantly larger (10% out of detected genes that were positively associated with astrocyte eigengene, N= 1,657) than the total number of genes that had higher expression in cocultured neurons (4% of all genes that were detected in both data sets, 610/13,968; p = 1.1×10^−22^, binomial test, **Figure 7D**). This suggested that a set of the same genes that associated with high astrocyte eigengene values were also induced in neurons with contact with glial cells. Out of these genes, 32 were synaptic (**Table S17**) and they were particularly enriched for members in the post-synapse specialization (N = 19 genes, GO:0099572, q=1.9×10^−8^) and included members especially in synapse organization, signaling and postsynaptic processes (**Figure 7C**). Altogether, these findings suggested that the induction of neuronal genetic programs that associated with high values of the astrocyte eigengene were in substantial part driven by physical contact between the neurons and the glial cells. They further indicated that the contact-driven intercommunication between neurons and glia was particularly relevant in regulating expression of members in the postsynaptic components.

### Glial cells induce genetic programs related to schizophrenia in neurons

Given that recent genetic discoveries have highlighted neuronally expressed genes with functions in synaptic biology in severe psychiatric illnesses, including schizophrenia^3,5,40,41^, we wondered whether these loci overlapped with those genes that were found to be induced in neurons by astrocytes. To address this question, we calculated gene-wise associations from gene-surrounding variants using summary statistics from a recent genome-wide association study (GWAS) for schizophrenia^5^ using MAGMA^42^. Using the gene-wise associations, we performed a gene-set analysis for the 1,970 genes that were induced by astrocytes in neurons. The analysis revealed significant association for variants nearby the astrocyte induced genes with schizophrenia (β = 0.22, p = 1.6×10^−12^) (**Figure 7E, Table S18)**. We further divided the genes into those that possessed a synaptic annotation in SynGO^26^ and non-synaptic genes. This revealed that the association to schizophrenia was stronger, but not exclusive, to genes with known synaptic annotations induced by astrocytes (β_synaptic_ = 0.41, p = 2.5×10^−7^; β_non-synaptic_ = 0.18, p = 4.8×10^−8^). The induced expression of genes relevant to schizophrenia by astrocytes in neurons suggests that astrocytes regulate neuronal functions that go awry in schizophrenia.

To assess the specificity of the association with schizophrenia, we repeated the analysis for three other central nervous system phenotypes with variable ages of onset: autism spectrum disorder (ASD)^43^, general intelligence^44^, and Alzheimer’s disease (AD)^45^. The 1,970 induced neuronal genes showed modest association for AD (β = 0.05, p = 0.012) and IQ (β = 0.07, p = 0.009), while there was no evidence for association with ASD (**Figure 7E**). As for schizophrenia, the association for intelligence was stronger for genes with synaptic annotations than for genes without such annotations (β_synaptic_ = 0.16, p = 0.02; β_non-synaptic_ = 0.05, p = 0.05). To investigate whether the 1970 induced neuronal genes were affected in patients with schizophrenia, we studied their overlap with genes that were previously identified reduced in the prefrontal cortex of schizophrenia patients^46^. Of 314 genes that had been found to be reduced in expression in patients, 50 were among the 1,970 genes we found to be induced by astrocytes in neurons (1.35-fold enrichment, p=0.028, binomial test). Together these results demonstrated that the presence of astrocytes in glial cultures was associated with dose-dependent induction of genetic programs relevant particularly for schizophrenia in the accompanying neurons including synaptic function and neuronal maturation.

### Astrocytic genes that induce synaptic programs in neurons are associated with schizophrenia

The potential role of astrocyte-neuron interactions in schizophrenia prompted us to investigate whether the genes whose high expression in glial cells was linked to synaptic programs in neurons were involved in schizophrenia. We reasoned that since glial cells are central in regulating many of the neuronal processes that have previously been implicated by genetic studies in schizophrenia^47^, disturbances in the glia-neuron regulatory interactions may also be relevant for the disease. A gene set analysis in MAGMA for the 123 glial genes whose high expression was linked to induction of synaptic programs found that these genes tended to harbor common variation that was associated with schizophrenia (β = 0.28, p = 0.0063) (**Figure 7F, Table S19**) and was strongly driven by the genes with known synaptic annotations in SynGO (β = 0.91, p = 0.0050)^26^.

To investigate individual genes underlying the gene-set association for schizophrenia, we ranked the 123 glia genes according to the MAGMA Z-statistic of gene-wise associations (**Figure S5E, Table S20)**. Among the genes with the strongest associations comprised synaptic genes including the cell adhesion molecules *LRRC4, LRRC4B* and *NRXN1* as well as the astrocytic *LRP4* involved in regulation of glutamate transmission^35^, and included genes in 10 genome-wide significant loci (*LRP4, AP3B2, KCNB1, CCDC39, PCDHB4, MSI2, B9D1, NALCN, SNX32*) in recent schizophrenia GWAS from The Psychiatric Genomics Consortium (PGC3)^5^. In addition, two genes had direct genetic evidence from rare variant associations (*NRXN1* and *MAGI2*). ^3^. Our results indicated that these genes’ expression in glial cells was associated with synaptic programs and neuronal induction of genes associated with schizophrenia. This further suggested that the neurobiological programs in neuron-glia interactions *in vitro* are relevant to schizophrenia biology.

## Discussion

Here, we studied how glial cells impact neuronal cell state and maturation by analyzing the co-expression of genes between the two cell populations in coculture using cross-species-resolved RNA-seq data. Our analysis revealed that genetic programs in glial cells that covary in abundance with astrocyte markers induce pre- and postsynaptic programs in neurons that associate with schizophrenia (**Figures 3 and 7**). We discovered that the neuronal synaptic gene-expression programs were associated with high expression of astrocytic synaptic cell adhesion molecules including neurexins (*Nrxn1*), neuroligins (*Nlgn1, Nlgn3*), and leucine rich repeat transmembrane proteins (*Lrrc4, Lrrc4b*) with direct evidence from both common and rare variant associations to schizophrenia^3,5^ (**Figures 5–7**). Our analysis revealed associations with additional synaptic genes expressed by astrocytes with functions related to synaptic vesicle cycle (Cadps, Nrxn1, Unc13B, and Prkcb), as well as glutamate receptor subunit Gria1 and synaptic potassium channels Kcnb1. Importantly, we found that the synaptic cell adhesion molecules were induced in astrocytes upon coculture with neurons, suggesting that the synaptic-adhesion abilities of astrocytes are induced by the presence of neurons. Moreover, the glial cells in coculture had higher expression of genes encoding functions in the glutamate recycling pathway, implying that the cocultured astrocytes are recruited to participate in the neuronal processes of the glutamatergic neurons. Finally, we saw that the differentiating neurons in coculture enhanced gene programs characteristic of astrocyte maturation in the early postnatal murine astrocytes. This is also consistent with previous work supporting that neurons patriciate in regulation of astrocytes^16^.

Genetic associations in schizophrenia are concentrated in genes that are highly expressed by neurons and cortical regions of the brain with roles in the synapse, transmission and differentiation^3–5^. This has focused much of the previous attention in the field on neurons. Besides neurons, astrocytes are active modulatory components of neural circuits that shape the structure and function of neuronal synapses and ultimately behavior through direct contacts and secreted factors ^48^. Indeed, together with the presynaptic and postsynaptic nerve terminals, the synaptically associated astrocytes compose a solid functional unit known as a tripartite synapse ^49^. Importantly, astrocytes express many gene products typically described as neuronal synaptic elements, including receptors for neurotransmitters, synaptic cell adhesion molecules, and components of synaptic-like vesicle cycle^29,30,34,50^ in which gene variants have been associated with risk for schizophrenia^3,5,51^.

We found that physical contact with astrocytes was required to induce neuronal synaptic gene expression **(Figure 7A–D**). This is in line with previous reports showing that contact between astrocytes and neurons is required for synapse formation^30,52^. Trans-synaptic cell adhesion molecules, such as neurexins (NRXN), neuroligins (NLGN) and leucine rich repeat transmembrane protein (LRRTM) provide a structural scaffold that holds synaptic terminals together and participate from early steps of synapse formation to regulating synaptic plasticity^53,54^.

Astrocytes have been recently demonstrated to express NRXN1 and NLGN1 implying that they could also be involved in fastening astrocyte processes to the synapse^29,30^. Importantly, deletion of NRXN1 in astrocytes has been reported to affect synapse function by impairing the maturation of silent synapses, AMPA-receptor recruitment, and long-term potentiation without affecting the number of synapses^29^. Postsynaptic LRRC4 and LRRC4B associate with postsynaptic density (PSD95) to regulate excitatory synapse formation by LRRC4 binding to presynaptic netrin-G2 (NTNG2) and LRRC4B binding to receptor tyrosine phosphatases LAR (PTPRF), PTPRS, and PTPRD to induce functional presynaptic differentiation in contacting neurites^55–57^. In a manner similar to transsynaptic binding of NRXN1 and NLGN1^58^, the heterophilic connections with LRRC4 and LRRC4B to their binding partners enable correct connectivity between presynaptic and postsynaptic terminals^55,56^. Here, we report that expression of these genes encoding for synaptic cell adhesion molecules in glial cells correlates with the expression of genes with roles in synaptic organization including structural trans-synaptic adhesion molecules in neurons (**Figure 3**). Importantly, we found that *NTNG2* and *PTPRS* that bind LRRC4 and LRRC4B, respectively, were induced in neurons in response to astrocytes. These findings provide further evidence that these molecules participate in glia-neuron interactions that may involve synaptic contacts. The expression of heterophilic synaptic cell adhesion molecules characteristic for presynaptic and postsynaptic nerve terminals in the glial cells suggest that they may be involved in positioning glial processes with connections to the two neuronal synaptic terminals.

Astrocytes are central in regulating many of the key neuronal functions, related to development, synaptogenesis, maturation, and synaptic transmission that are fundamental to all information processing in the brain^6^. Together with the discoveries of synaptic gene variants in schizophrenia there has been increasing interest in astrocytes’ potential role in the disorder^47,59^.

Complement proteins are expressed by neurons and glial cells including astrocytes and are localized to a subset of developing synapses^60,61^. Defects in developmental pruning can lead to synaptic loss long before onset of clinical symptoms in line with the developmental model. Here we show that early developmental glia interactions with hPSC derived neurons with resemblance to late prenatal stages^15^ are associated with coregulated expression of synaptic adhesion genes linked with schizophrenia.

Follow up studies of recent genetic discoveries comparing gene expression patterns across tissues and cell types have implicated neurons as the cell type in which schizophrenia-implicated genes tend to have strongest expression on average^3–5^. However, many emblematic neuronal functions depend on interplay with glial cells and may impact the pathology through direct or cell-nonautonomous effects on neurons. Here we show that expression of schizophrenia-associated genes in glial cells is correlated with a neuronal maturation state and induction of the expression of risk genes in neurons. Our results underscore the importance of evaluating the converging functional impact of emerging genetic discoveries in living biological model systems that provide mechanistic insight, such as cellular interactions, into affected biological processes.

## Supporting information

Supplemental tales S1-S20

## Limitations of the study

Here we showed that glial cells induce synaptic gene-expression programs in iPSC-neurons in coculture, and that these programs are enriched for genes implicated in schizophrenia. We found that the induction of these programs in neurons was in turn correlated with the expression of diseased-linked synaptic cell adhesion molecules in glial cells whose expression was dependent on neuronal contact with glial cells. Although we show that synaptic gene programs in neurons are correlated with the expression of pro-synaptic genes in glia, it remains unknown how these interactions are mediated. We have conducted a series of experiments in mixed cultures of human induced neurons and mouse glial cells that revealed a potential relevant cellular context for schizophrenia. Although mouse glial cells have been shown to enhance neuronal maturation, it is possible that they lack abilities that are inherent for human cell types and will only be present in fully human coculture systems. As human astrocyte and human neuron co-culture paradigms are developed and optimized, it will be important to test to which extent the cellular programs we report here are conserved. Overall, the *in vitro* cell cultures are a flexible experimental platform to study cellular responses to their surroundings. However, although powerful, this approach does not provide a comprehensive view of the full complexity of the true *in vivo* cellular context in the brain. Instead, it aims to illuminate individual components of neuronal interactions with other cell types. Finally, schizophrenia is a behavioral human specific phenotype with both genetic and environmental influences^62,63^. It is therefore, important to note that we do not model schizophrenia but cellular processes underlying the disorder.

## Author contributions

The project was conceived and designed by OP, RN, and KE. AT, MT, HG and TC performed the experiments, with supervision from RN. JM performed the village experiment, with supervision from SAM and KE, and analysis by DM. VV performed the immunostaining and imaging, using equipment in SLF’s lab. All bulk RNAseq analyses were conceived, designed, and performed by OP. Single cell RNAseq data was processed and analyzed by DM, EV, in supervision of SM and OP. OP and RN wrote the initial draft of the manuscript and figures. All authors contributed to editing of the manuscript.

## Acknowledgements

We would like to warmly thank the voluntary participants who have donated the cell lines for research. This work was funded by U01MH105669 (NIH/NIMH) and the Stanley Center for Psychiatric Research at the Broad Institute, with additional support from R37NS083524 and U01MH115727. RN was also supported by a NARSAD young investigator award (Brain and Behavior Research Foundation). OP was additionally supported by Instrumentarium Science Foundation, Sigrid Juselius Foundation, Orion Research Foundation, Jalmari and Rauha Ahokas Foundation, Päivikki and Sakari Sohlberg Foundation, Jenny and Antti Wihuri foundation, and Niilo Helander Foundation.

---

**Figure S1.**
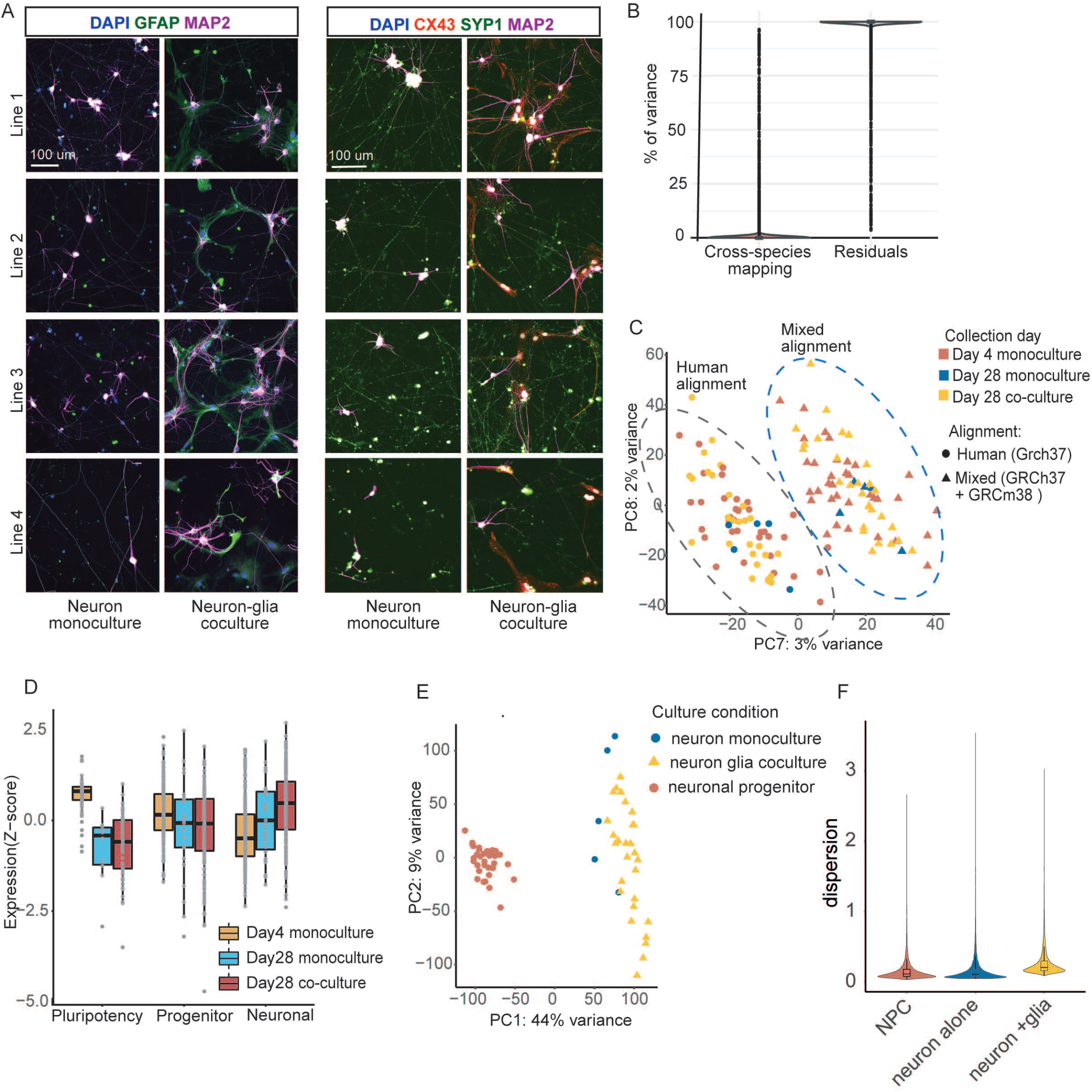
Alignment effect on transcriptome. A) Representative images of neuron monocultures and neuron-glia co-cultures from 4 independent cell lines, staining for left: DAPI (blue) GFAP (green) and MAP2 (magenta), and right: DAPI (blue) CX43 (red) Syp1 (green) and MAP2 (magenta), Scale bar= 100um. B) Variance partitioning by human and mixed reference genome reveals 329 genes for which cross-species mapping explains >10% of variation in read counts. On average cross-species mapping explains only 0.78% of the variation in the read counts. C) PCA components 7 and 8 the cluster data set acording to alignment. D) PCA of RNAseq data from day 28 neurons aligned to mixed reference genome. E) Canonical marker gene expression for pluripotency, neuronal progenitor cells and neuronal identity at different stages of differentiation and culture conditions.

**Figure S2.**
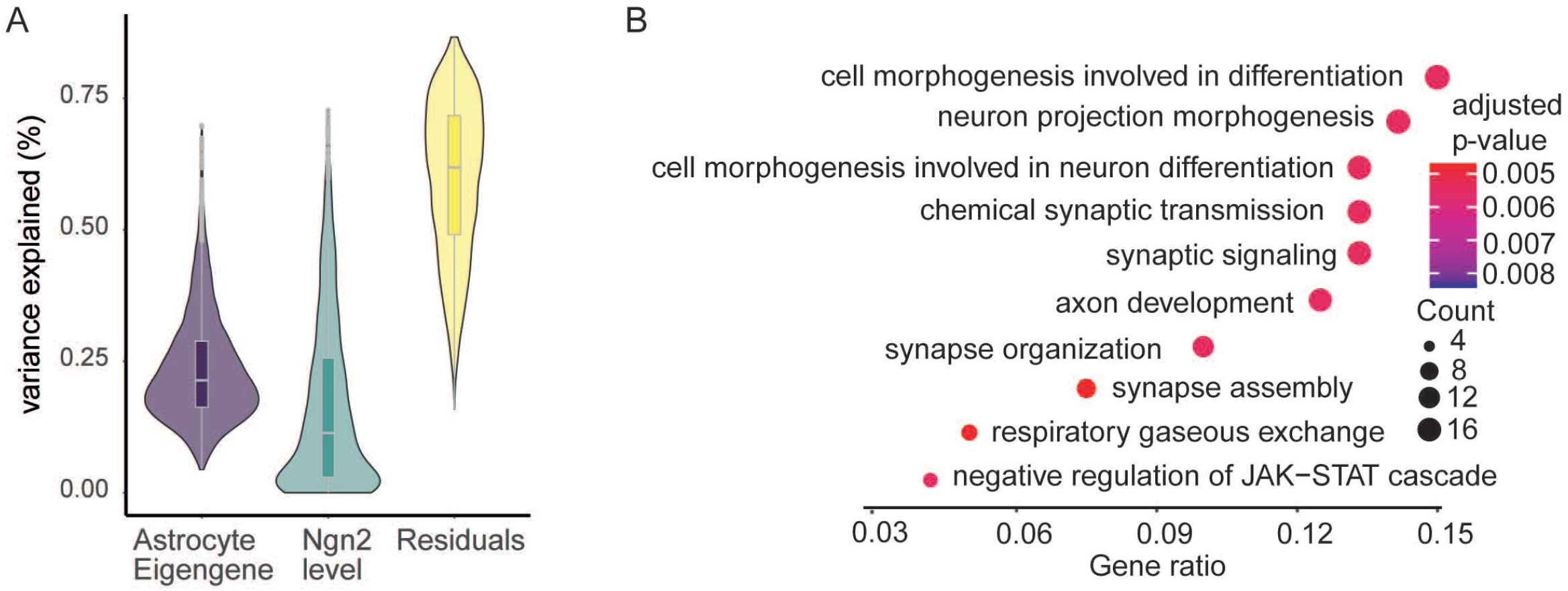
Variance explained by model covariates and GO terms for genes induced by astrocytes in neurons. A) Variance explained in gene expression for model covariates. B) Transcriptome-wide significantly induced genes are enriched for functions in neuronal development and synapse

**Figure S3.**
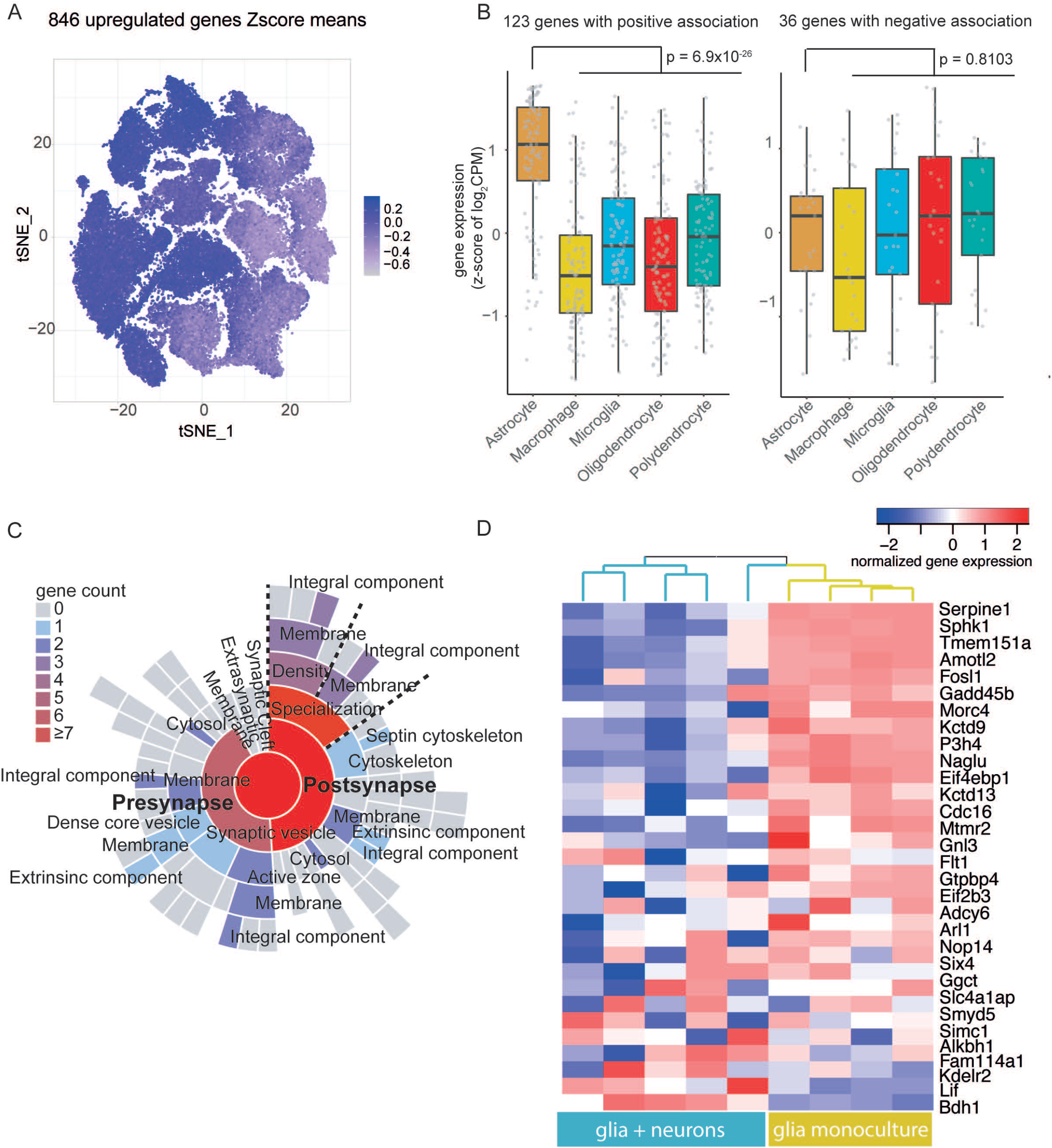
Glial genes that associate with neuronal maturation are predominantly expressed by mouse astrocytes and enriched for synaptic genes. A) tSNE projection of scRNA-seq data showing expression of genes induced in coculture. B) The 123 positively associated genes had the highest expression in astrocytes in the single cell atlas of adult mouse brain. The 36 negatively associated genes were uniformly expressed across glial cell types in the mouse brain atlas. C) The 159 glial genes have synaptic functions in SynGO. D) Glial genes with negative association to neuron maturation are reduced in astrocytes after contact with neurons.

**Figure S4.**
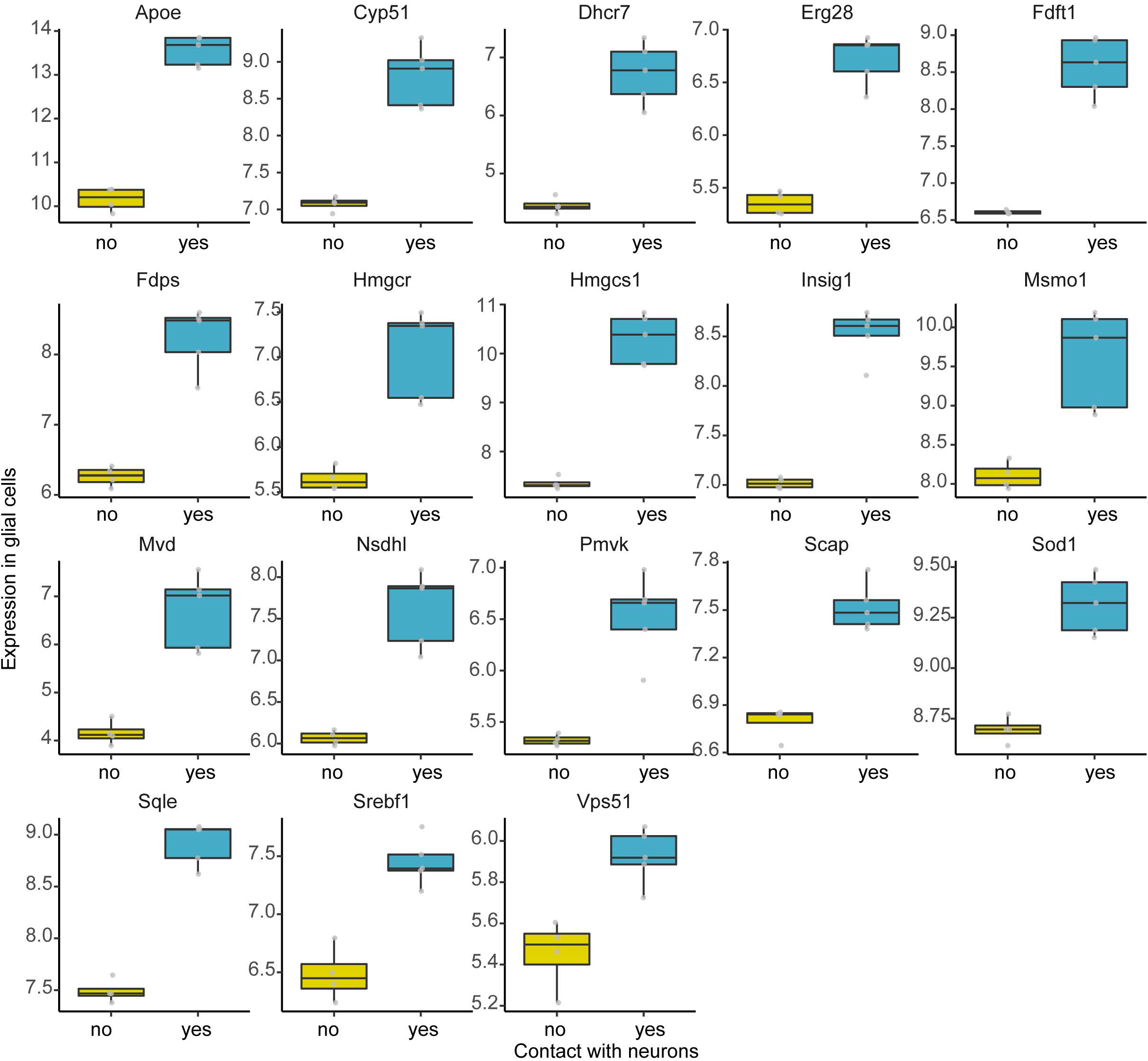
Transcripts encoding members of the cholesterol synthesis pathway are induced in astrocytes after contact with neurons.

**Figure S5.**
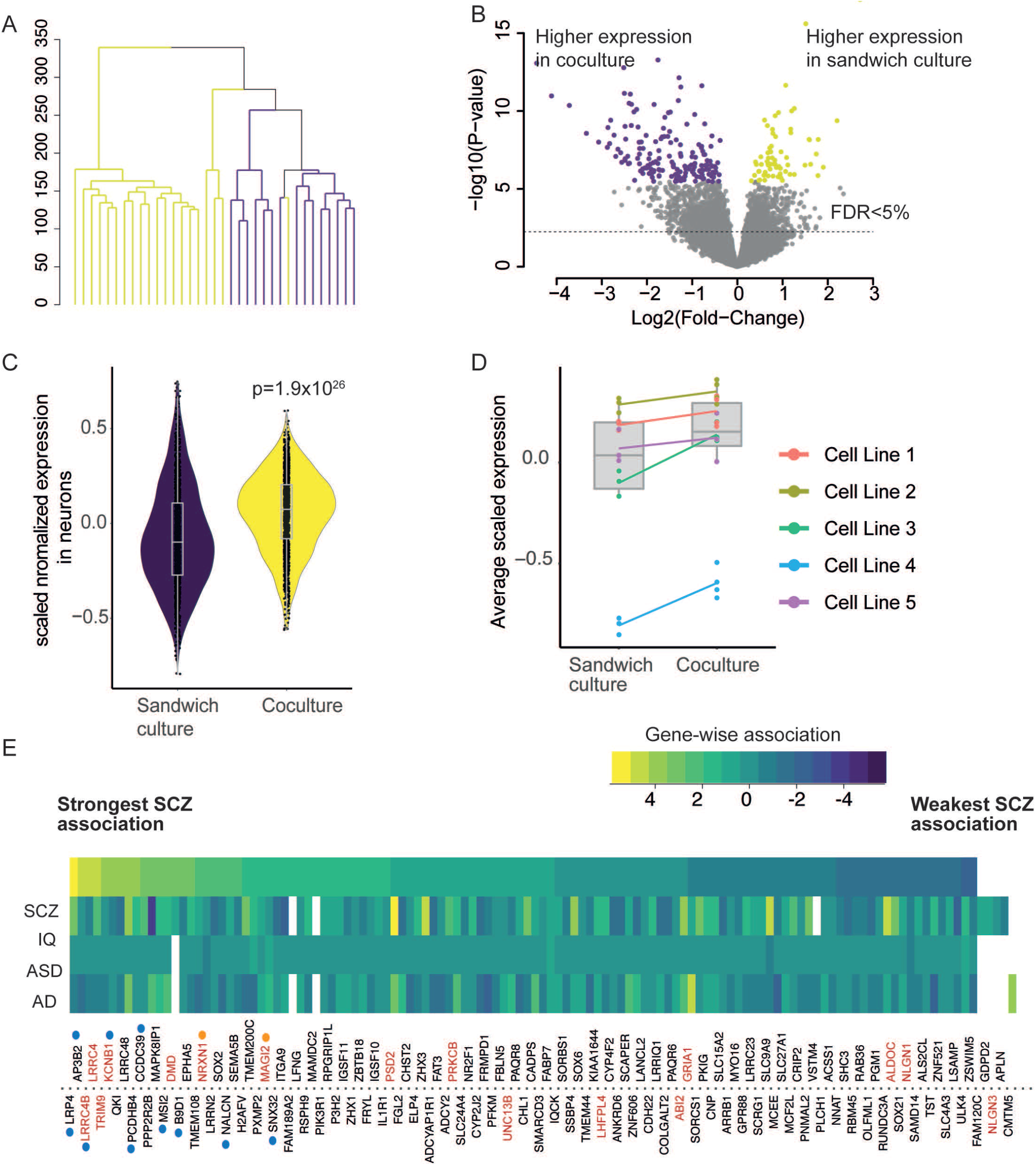
Neurons in sandwich culture and coculture with glial cells have global transcriptional changes. A) Hierarchical clustering of neurons in sandwich and coculture. B) Volcano plot of differential gene expression between neurons in coculture and sandwich culture.C) Neurons in coculture have on average higher expression of genes that associate with the astrocyte eigengene (p=1.9×10^26^ paired t-test). D) The average levels of the transcripts that associate with the astrocyte eigengene are consistently higher in all neuronal lines. E) Gene-wise associations for 123 glial genes whose high expression induce synaptic gene programs in neurons. The heat-map coloring indicates the standardized z-score of gene-wise p-values from MAGMA. The genes are ordered by the strength of association (z-score value) to schizophrenia. Synaptic genes are colored in red. Genes with direct evidence through coding variants and fine mapping in schizophrenia are indicated by arrows. Genes located in genome-wide significant loci in PGC3 for schizophrenia are highlighted by blue boxes.

