## Supplemental methods for "Astrocytic cell adhesion genes linked to schizophrenia correlate with synaptic programs in neurons"

### Materials and Methods

#### *Human pluripotent stem cell line cohorts and derivation*

Human induced pluripotent stem cell lines were derived from a multi-institutional collection of fibroblasts and peripheral blood mononuclear cells (PBMC) lines from population controls originating from the Swedish Schizophrenia Cohort (Karolinska Institute), the Northern Finnish Intellectual Disability Cohort (NFID) (Kurki et al., 2019), Umea University, Massachusetts General Hospital (MGH), McLean Hospital, and GTEx. The iPSC lines were reprogrammed in house (Nehme et al., 2018), at the New York Stem Cell Foundation (NYSCF) or at the Harvard Stem Cell Institute (HSCI) iPS core as described previously (Nehme et al., 2022) (**Table S1**). The discovery cohort additionally included three embryonic stem cell lines from the Human Embryonic Stem Cell Facility of the Harvard Stem Cell Institute.

#### hPSC culture

Human ESCs and iPSCs were maintained on plates coated with geltrex (life technologies, A1413301) in StemFlex™ medium (Gibco™, A3349401) supplemented with Normocin™ Antimicrobial Reagent (Invivogen, Ant-nr-1). and passaged with accutase (Gibco, A11105). All cell cultures were maintained at 37°C, 5% CO2.

#### Infection of hPSCs with lentiviruses

Lentivirus particles were produced by Alstem (http://www.alstembio.com/). hPSCs were seeded in a geltrex coated 12 well plate at a density of 100,000 cells/cm$^2$ in StemFlex medium supplemented with rock inhibitor (Y27632, Stemgent 04-0012) and lentiviruses containing a murine Neurogenin 2 (Ngn2) tagged with a puromycin resistance gene and a tetracycline inducible GFP, at a MOI (multiplicity of infection) of 2. Cells were plated at a density of

100,000 cells/cm2 and incubated while in suspension with media containing 10 uM ROCK-Inhibitor (Sigma, Y27632). After 24 hours, the medium was changed to StemFlex. The cells were grown until confluency, and then either maintained as stem cells, passaged, banked, or induced with Doxycycline for neuronal differentiation.

**Neuronal differentiation**

Neuronal differentiation of PSCs into cortical glutamatergic neurons was carried out as previously described (Nehme et al., 2018). In brief, the differentiation was carried out by adding Doxycycline hyclate (2 μg/mL) to N2 supplemented media (Thermo Fisher, 17502048,) with patterning factors SB431542 (Tocris, 1614, 10 μM), XAV939 (Stemgent, 04-00046, 2 μM) and LDN-193189 (Stemgent, 04-0074, 100 nM), as described previously(Busskamp et al., 2014; Nehme et al., 2018; Zhang et al., 2013). Puromycin selection was used (5μg/μL), from days 2 to 6 to remove non-transduced cells. At 4 days post induction, neuronal cells were resuspended into Neurobasal media (Gibco, 21103049) that was supplemented with B27 (Gibco, 17504044, 50X), doxycycline (2 μg/mL), brain-derived neurotrophic factor (BDNF), ciliary neurotrophic factor (CTNF), and glial cell-derived neurotrophic factor (GDNF) (R&D Systems 248-BD/CF, 257-33 NT/CF, and 212-GD/CF at 10 ng/mL each). From this point onwards the neurons were either co-cultured with murine glial cells that were derived from early postnatal (P1-P3) mouse brains as described previously (Di Giorgio et al., 2008) or were left to grow as monocultures (mouse strain https://www.jax.org/strain/100012; animal ethical committee approval by Harvard University: Animal Experimentation Protocol (AEP) # 93-15). To evaluate whether physical contact between neurons and glia was required for some of the transcriptional effects, we used a sandwich culture set-up whereby glia cells were cultured at the bottom as a monolayer, and neuronal cells were grown on a membrane insert (Sigma, CLS3450).

**Cell villages**

The cell villages were generated as described previously (Mitchell et al., 2020; Wells et al., 2021). At day 3 of the differentiation, the differentiating cells from 48 donors were passaged into differentiation media that was supplemented with 5-Ethynyl-2'-deoxyuridine (Life Technologies, A10044, 10 µM). Cell villages of the immature neurons were generated at day 6 by dissociating the cells with Accutase®, counting them by the Scepter™ Automated Cell Counter (Millipore Sigma) and plating the cells at a density of 40 000 cells/cm$_2$ in Neurobasal media (Gibco, 21103049) supplemented with B27 (Gibco, 17504044, 50X), doxycycline (2 µg/mL), brain-derived neurotrophic factor (BDNF), ciliary neurotrophic factor (CTNF), and glial cell-derived neurotrophic factor (GDNF) (R&D Systems 248-BD/CF, 257-33 NT/CF, and 212-GD/CF at 10 ng/mL each). Neuronal villages were either grown as monocultures or co-cultured with murine glial cells (at a density of 70 000 cells/cm$_2$). Villages were harvested for single cell RNA sequencing at day 28 of the differentiation.

**Immunohistochemistry**

Cultures neurons were fixed at D28 of neuronal differentiation in 4% paraformaldehyde + 5% sucrose in DPBS for 20 min at room temperature. Cells were incubated with blocking buffer containing 4% horse serum, 0.1M Glycine, and 0.3% Triton-X in PBS for 1 hour at room temperature. Primary antibodies, diluted in 4% horse serum in PBS, were incubated overnight at 4oC. Secondary antibodies were diluted in 4% horse serum in PBS and applied for 1 hour at room temperature. Samples were washed 3x with PBS and imaged on spinning disc confocal microscope (Andor Dragonfly) with a 20x air interface objective using Fusion software. The following antibodies were used: chicken anti-MAP2 (1:10,000, Abcam ab5392), Rabbit anti-Syp-1 (1:1000 millipore AB1543P), Rabbit anti-GFAP (1:250, ab16997), Rabbit anti-Cx43

(1:500, ab230537). Goat Alexafluor plus-555- conjugated anti-mouse (Cat # A32727) and Goat anti-Rabbit Alexafluor plus-488 (Cat # A32731)

**RNA sequencing**

For bulk RNA sequencing cells were harvested in RTLplus Lysis buffer (Qiagen 1053393) and stored at -80°C. Each experiment was conducted in 3-4 replicates to reduce experimental variability. Sequencing libraries were generated from 100 ng of total RNA using the TruSeq RNA Sample Preparation kit (Illumina RS-122-2303) and quantified using the Qubit fluorometer (Life Technologies) following the manufacturer's instructions. Libraries were then pooled and sequenced by high output run on a HiSeq 2500 (Illumina). Cells grown using the sandwich culture set up (along with neuron monocultures from the same cell lines and glia monocultures from the same batch) were processed and sequenced at the Broad Genomics Platform using the Smartseq2 workflow (Picelli et al., 2014). The RNA-seq fastq data was aligned to ENSEMBL human reference genome (build GRCh37.p13/hg19) and mixed human-mouse reference genome (GRCh37/hg19 and GRCm38/*mm10*, GSE63269) (Macosko et al., 2015) by STAR (v.2.5) (Dobin et al., 2013). Prior to genome aligning, the Illumina adapters and low-quality base-pairs were clipped from the ends of the sequence reads by Trimmomatic (v.0.36) (Bolger et al., 2014) and reads with length < 36 base-pairs were removed. The gene-wise read-counts were generated from the aligned reads by featureCounts in Rsubread (v.1.32) (Liao et al., 2014) using GENCODE GTF annotation version 19 and a custom GTF file for the mixed genome (GSE63269)(Macosko et al., 2015) for the mixed genome. The reads from the three experimental replicates were summed together yielding an average library size of ($11\times10^6$ reads). The final analyses were carried out from read counts from the mixed alignment.

For single cell RNA sequencing of neuron villages, the cells were harvested, and RNA samples prepared with 10X Chromium Single Cell 3' Reagents V3 followed by sequencing with

Illumina NovaSeq 6000 (Illumina) using a S2 flow cell at 2 x 100bp, as previously described (Limone et al., 2022; Wells et al., 2021). Raw sequence fastq- files were then demultiplexed and aligned according to the Drop-seq workflow (Macosko et al., 2015) to human reference genome (GRCh38, ensembl v89 gene model), and filtered for high quality mapped reads (MQ>10). Donor reference genotype data was processed as previously described and aligned to human reference (GRCh38) (Mitchell et al., 2020) The Dropulation algorithm was then run on the preprocessed sequence data and VCF data using strict quality control criteria for variant filtering to accurately specify the donor identity of each droplet, as specified previously (Wells et al., 2021). The Dropualtion algorithm analyses each cell line data independently and generates a likelihood of the data for having been generated by each of the donors that are included int the VCF file. The donor identity is assigned as the computed diploid likelihood at each UMI summed up across all sites. (Wells et al., 2021). For gene expression analysis, the digital gene expression matrices were generated, and the UMI counts of all assignable single cells per donor were summed to generate a donor by gene matrix. The differential expression analysis was performed with voom-limma while adjusting for covariates.

**Cross-species mapping**

The gene-wise cross-species mapping effect was estimated by variancePartition package (v.1.8.1) in R, which fits mixed linear model to estimate sources of variation in the data (Hoffman and Schadt, 2016). The analysis was run on $\log_2$-transformed normalized human reads from data aligned to the two reference genomes. The alignment was included as a random effect in the model. The global effect of cross-species mapping was estimate by PCA on $\log_2$-transformed normalized RNAseq data from the two alignments.

**Differential gene expression**

Differential expression analysis and bulk RNAseq data processing was carried out separately for human and mouse, with the exception of mouse Ngn2 that was processed together with human data. Only data from day 28 of differentiation was included and separately processed for differential expression analysis. The differential gene expression was analyzed in limma-voom (v. 3.34.9) package (Law et al., 2014) using read counts from the mixed-genome alignment. The raw read counts were normalized for library size (method = TMM) and by voom to produce $log_2$-transformed normalized gene expression estimates. Genes with low read counts (> 10 reads in at least one library and total read count >15) were removed by filterByExpr function in edgeR (v. 3.20.9) prior to normalization. For differential expression analysis of astrocyte eigengene effect on neuronal gene expression a multifactorial model was used with astrocyte eigengene and normalized Ngn2 expression as covariates (~ AstroE + ngn2). The proportion of variance explained by each covariate was estimated with variancePartition package (v.1.8.1). For differential expression analysis of mouse RNAseq data a linear model with neuron eigengene as a covariate was used (~NeuroE). The p-values were adjusted with FDR < 5% and Bonferroni correction for transcriptome-wide significance. PCA was performed on normalized $log_2$ transformed CPMs to assess the global transcriptome. For PCA RNAseq data from day 4 NPCs was included and the data set was processed together with the neurons. Combat function of the SVA package (v. 3.26.0) was used to correct for experimental batches prior to PCA. For analyzing the effect of physical contact between glial cells and neurons bulk RNAseq data from isogenic experiments of neurons differentiated in glia coculture or in sandwich culture was analyzed. A PCA of the transcriptomic data was performed after adjusting for unwanted experimental variation using Combat. The differential expression analysis was done by limma-voom for human reads and adjusting for cell line effect, as well as sequence library quality measures from Picard tools (PCT_USABLE_BASES and MEDIAN_CV_COVERAGE) that were found to differ significantly between the isogenic experiments. The impact of neuron presence on glial cells was estimated by differential expression analysis of mouse genes in limma-voom of 4 replicates of glia alone cultures and 5 glia-neuron cocultures that were processed together. The experimental replicates (4 from each donor) from the cocultured glial cells were summed together to match the glia library size of the four replicates from glia alone cultures. The raw reads were then filtered (> 10 reads in at least one library and total read count >15) and normalized byy TMM-method in limma-voom package. The differential expression analysis was done between the two culture condition (~0 + culture.condtion). For analyzing the effect of glia coculture on neurons, UMIs of all donor cells in the two culture conditions were summed to generate meta cells. The differential expression analysis weas carried out for human genes in limma-voom between neurons grown with or without glial cells. The analysis was adjusted for donor sex, average number of UMIs per cell and number of cells of each donor as covariates. The donor effect was adjusted for using the block design in limma.

**Marker gene expression and eigengene generation**

Expression of canonical marker genes from previous literature were used to characterize neuron and glial cell populations in the different experiments (**Table S2**). Singular value decomposition was used to generate astrocyte and neuron eigengenes from $log_2$ normalized read counts per million (CPMs) using svd function in R. An m × n matrix of genes (m) and cell lines (n) for six canonical astrocyte marker genes selected from literature were used to generate the astrocyte eigengene (the first right singular vector v). The average variance explained by the eigengene was calculated from the squared singular value (d) of each gene divided by their sum $\frac{d^2}{\sum d^2}$ . The astrocyte eigengene values for the joint analysis of the discovery and deletion lines was calculated jointly. The eigengene value for glia-free cultures were set to zero. For neuron eigengene, 1,970 genes that were found positively associated with astrocytes were used to calculate the first singular vector (v).

#### *Gene set enrichment analysis*

GO term enrichment was analyzed using enrichGO function in clusterProfiler package (v. 3.8.1) in R with org.Hs.eg.db and org.Mm.eg.db (v. 3.6.0.) for human and mouse genes, respectively, using ENSEMBL identifiers. A minimum of 10 and maximum of 1000 genes per category were included to the analysis. P-values for adjusted for multiple testing by Benjamini & Hochberg method with p-value < 0.01 and q-value < 0.05 cutoffs. A custom gene universe was used as the background for the enrichment analysis including only genes that were analyzed in the RNAseq data. The GO terms for each category were analyzed separately. For synaptic gene annotations we used manually curated annotations from the SynGO database (Koopmans et al., 2019). Enrichment analysis with Synaptic GO terms were analyzed in the SynGO online portal (v.1.0) using custom gene universe of genes that were included in RNAseq data analysis (www.syngoportal.org ).

#### Mouse Brain single Cell Atlas

A single cell gene expression atlas from adult mouse brain was used to analyze the expression of glial genes *in vivo* (Saunders et al., 2018). Meta cells for available non-neuronal cell types including polydendrocytes, microglia, astrocyte, oligodendrocyte and macrophacge were downloaded and processed in limma-voom. Astrocyte meta cell was formed as the average of Gja1 positive and Gja1/Myoc positive meta cells. The gene expression in the glial meta cells was standardized to z-score values and t-test was used to analyze differences in average expression between the meta cells.

**Gene set association analysis**

Summary statistics from recent well-powered GWAS studies for four central nervous system traits, including schizophrenia (Trubetskoy et al., 2022), ASD (Autism Spectrum Disorders Working Group of The Psychiatric Genomics, 2017), measure of general intelligence (Savage et al., 2018), and AD (Jansen et al., 2019) were used to calculate gene-wise association in MAGMA (v.1.07b) (de Leeuw et al., 2015). SNPs from GWAS summary statistics were annotated to genes using 10kb window (window = 10) and gene locations from human genome build GRCh37.3 provided on the software website. The gene-wise associations were then calculated based on SNP annotations using 1000 genomes references data of European ancestry (1000 Genomes Project et al., 2015) for estimating the LD structure. Gene-set associations were analyzed from gene-wise associations with --gene-set flag for specified gene sets from the differential expression analysis. For mouse genes, human orthologues were identified prior to the analysis with getLDS function in biomaRt package (v. 2.34.2) using BioMart ENSEMBL databases for mouse and human.
